# Environmental chemical mixtures reprogram mammary epithelial development to epigenetic states associated with breast cancer

**DOI:** 10.64898/2026.04.01.715968

**Authors:** Meadow Parrish, Meloryn Seraj, Helia Nikoueian, Nicole Traugh, Andrew Chen, Piyush B. Gupta, Stefano Monti, Charlotte Kuperwasser

## Abstract

Environmental exposures occur as complex mixtures, yet the mechanisms by which they alter human tissue development and confer cancer susceptibility remain poorly defined. Here, we establish a physiological 3D human breast organoid high-content screening (3D-HCS) platform that quantitatively links exposure-induced developmental disruption to cancer-relevant cell states. Screening of structurally diverse environmental chemicals as single agents but also as mixtures reveals heterogeneous but reproducible developmental responses, indicating that distinct environmental chemicals perturb different aspects of mammary morphogenesis. Focusing on bisphenols, we show that a physiological-dose mixture (BPA, BPS, and BPF; 3BPX) disrupts organoid development and induces transcriptional programs characterized by extracellular matrix remodeling, epithelial plasticity, and partial epithelial–mesenchymal transition (EMT), accompanied by widespread DNA methylation remodeling. The 3BPX exposure signature maps to ER⁺ luminal breast cancers and is preferentially enriched in invasive lobular carcinoma (ILC), a subtype characterized by epithelial plasticity and stromal interaction. Genome-wide DNA methylation profiling revealed that the 3BPX-associated methylation signature is detectable in primary tumors and adjacent normal tissue and is enriched in Normal-like tumors and ILC. Together, these findings demonstrate that environmentally relevant chemical exposures induce heterogeneous developmental perturbations that converge on conserved transcriptional and epigenetic programs associated with epithelial plasticity, tissue remodeling, and cancer susceptibility. These exposure-induced states persist as molecular “scars” that map to defined epithelial and stromal compartments in human breast cancers, supporting a model in which developmental exposures establish a field of increased oncogenic potential.

**Summary Statement:** Using a human breast organoid high-content screening platform, we show that real-world chemical mixtures induce heterogeneous developmental disruption that converges on conserved EMT, wound-response, and epigenetic programs. These exposure-induced states persist as molecular “scars” and map to specific epithelial and stromal compartments in human breast cancers, particularly invasive lobular carcinoma, suggesting a developmental origin of cancer susceptibility.

## Introduction

Endocrine-disrupting chemicals (EDCs) are pervasive contaminants in air, water, food, and consumer products, and human exposure to these agents begins in utero and continues throughout life ^1–3^. National Health and Nutrition Examination Study (NHANES) biomonitoring shows that >95% of Americans carry detectable levels of multiple endocrine-disrupting chemicals (EDCs), including PFAS, bisphenols, phthalates, and parabens, often from birth^3,4^. EDC exposure is therefore ubiquitous, continuous, and multi-chemical, meeting criteria for population-level cancer drivers^5–8^. A growing body of epidemiologic and experimental evidence suggests that developmental exposure to EDCs may contribute to the rising incidence of hormone-sensitive cancers, including breast cancer, in young women ^6,8–12^. Of particular concern is exposure during critical windows of mammary gland development, when the breast epithelium is highly plastic and uniquely sensitive to environmental cues^6,9^. Yet, despite decades of concern, the mechanistic pathways through which EDCs perturb human mammary morphogenesis or increase carcinogenic susceptibility remain poorly defined.

To model real-world exposure scenarios, we selected a panel of environmental chemicals representing distinct but relevant biological classes (Supplemental Table 1). These include EDCs, such as bisphenols, parabens, and phytoestrogens (eg. S-equol), which are widely detected in human populations and are known to modulate hormone receptor signaling^15–17^. We also included compounds such as PFAS and PFOS, which exhibit endocrine and metabolic activity^18^, and kynurenine, a bioactive metabolite implicated in immune and metabolic regulation^19^. In addition, we incorporated benzo[a]pyrene (BaP), a well-characterized environmental carcinogen, to represent genotoxic exposures that may occur concurrently with EDCs^20^.

Most insights of chemical exposures are derived from rodent models or in vitro systems that incompletely capture the architecture, lineage diversity, and developmental programs of the human breast^13,14^. Recent advancements in human organoid technology offer an opportunity to overcome limitations in 2D and animal models. Three-dimensional (3D) organoids generated from primary human breast tissue recapitulate key features of mammary development, including progenitor hierarchies, branching morphogenesis, and differentiation programs^22^. We recently demonstrated that when coupled with multimodal imaging and computational analysis, organoid systems can serve as powerful platforms for studying microenvironmental effects on human tissue development ^23–25^. However, scalable platforms that can integrate high-content morphological profiling with molecular and epigenetic readouts under real-world exposure conditions remain lacking.

To address this challenge, we developed an AI-assisted 3D human breast organoid high-content screening (3D-HCS) platform capable of quantitatively measuring developmental disruption and morphogenic alterations across organoids exposed to chemicals. This approach allowed us to compare how distinct classes of environmental agents, including both non-genotoxic signaling modulators and DNA-damaging agents, influence human mammary development individually and in combination. We reveal that continuous bisphenol exposure modifies the trajectory of organoid development and responsiveness to genotoxic insults. Moreover, we also identify conserved, exposure-induced transcriptional signatures as well as DNA methylation “scars” that mirror patterns detectable in human breast cancers, suggesting that early-life or developmental exposures may leave durable molecular fingerprints that prime breast tissue for future oncogenic transformation.

## Results

### 3D High-Content Screening of Human Breast Organoids Identifies Heterogeneous Developmental Responses to Environmental Chemicals

We recently developed and characterized a 3D human breast hydrogel organoid model that recapitulates the key stages of embryonic mammary tissue organogenesis including stem cell induction, patterning, morphogenesis (ductal initiation, branching morphogenesis), and alveolar maturation ^24,25^. Organoids seeded from single human donor epithelial cells in this model form terminal ductal-lobular unit structures (TDLU) by 21 days when embedded into a defined 3D hydrogel matrix composed of Type I Collagen, hyaluronic acid, laminin, and fibronectin (Supplemental Figure 1a). Notably, in this model, organoids establish a mesenchyme-like stroma that emerges from forming organoids through an epithelial-to-mesenchymal transition (EMT). This mixed parenchyma/mesenchyme organoid model establishes epithelial–stromal interactions that mirrors the crosstalk during development and branching morphogenesis^24^.

Using this platform, we conducted a confocal 3D high-content screen (3D-HCS) to assess how continuous exposure to environmental chemicals might perturb human mammary organogenesis, development, and morphogenesis in a model that closely mirrors breast development (Fig. 1A). Single human breast epithelial cells were seeded into 3D hydrogels and exposed continuously to various chemicals for 21 days at a 100nM dose (Supplemental Table 1). These chemicals were selected to represent a range of environmental exposures with diverse mechanisms of action, including endocrine signaling modulators, metabolic regulators, and compounds with known or suspected carcinogenic activity (Supplemental Table 1). 3D hydrogel cultures were allowed to develop into organoids and brightfield images were captured using confocal microscopy. An automated and AI-assisted segmentation pipeline was developed to identify epithelial contours and branching architecture in order to quantify organoids from Maximum Intensity Projections (MaxIP) of Z-stacks (Supplemental Figure 1b). This approach enabled automated quantification of 1) total organoid number (Fig. 1b), 2) organoid size distribution (Fig. 1c,d), and 3), organoid complexity (Fig. 1e,f) across 4 different hydrogel replicates for each chemical treatment.

**Figure 1.**
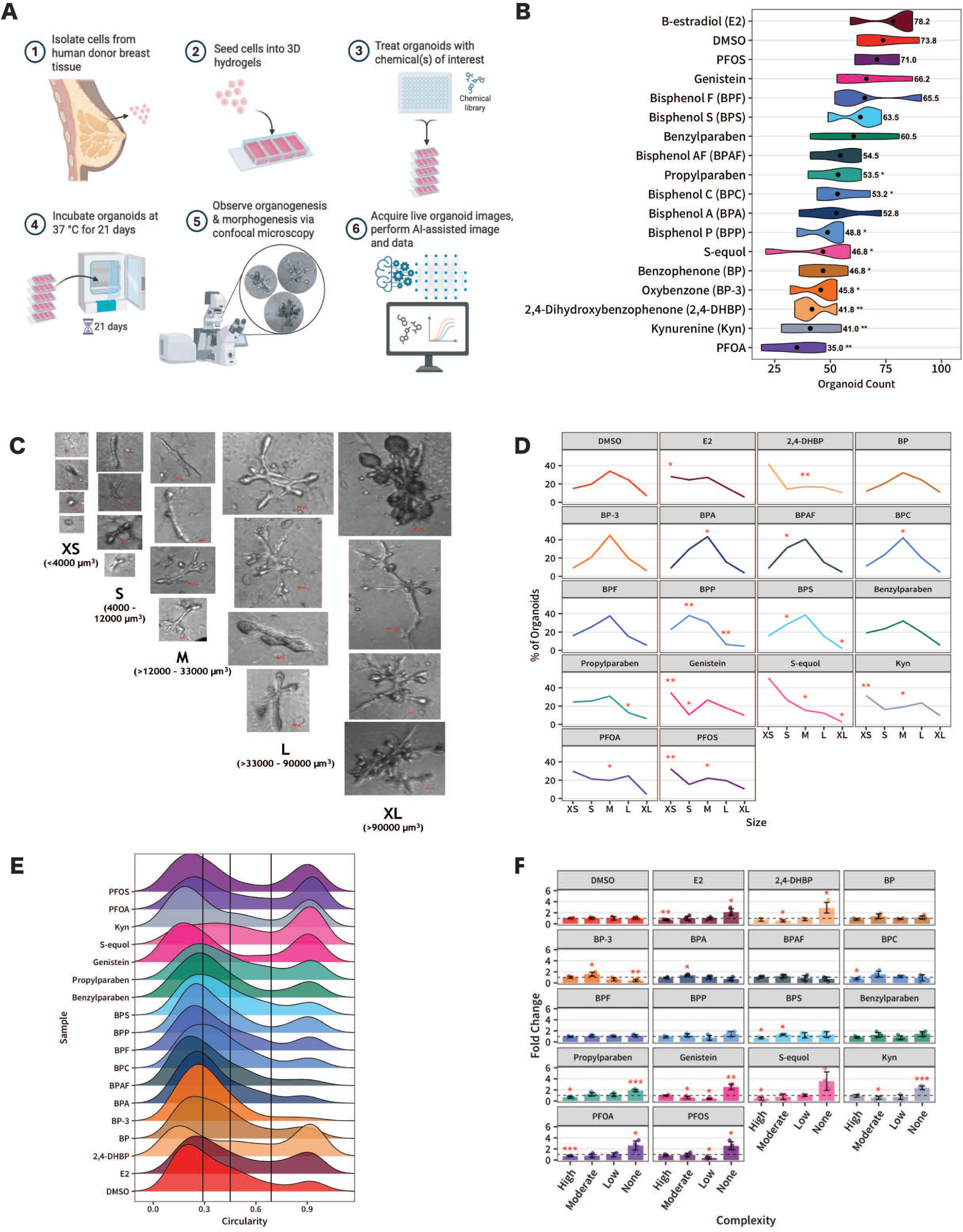
High-content screening (HCS) platform reveals exposure-dependent alterations in human breast organoid morphology. **(A)** Schematic of the 3D high-content screening workflow. Primary epithelial cells isolated from human donor breast tissue were embedded in 3D hydrogels, allowed to undergo organoid morphogenesis, and treated with endocrine-disrupting chemicals (EDCs) or vehicle control (DMSO). Organoids were cultured for 21 days and imaged by confocal microscopy, followed by automated image acquisition and quantitative morphometric analysis. **(B)** Organoid counts per condition (n = 4 biological replicates per treatment). Black dots indicate mean values. Statistical significance relative to DMSO was assessed using unpaired Welch’s *t*-tests (*p* < 0.05, \**p* < 0.01). **(C)** Representative bright-field images of organoids stratified by size category—extra-small (XS), small (S), medium (M), large (L), and extra-large (XL)—based on projected area. Scale bar, 100 µm. **(D)** Distribution of organoid sizes across treatment conditions, expressed as the percentage of organoids within each size bin. Statistical significance relative to DMSO was determined using unpaired Welch’s *t*-tests (*p* < 0.05, \**p* < 0.01). **(E)** Ridgeline plots showing the distribution of organoid circularity across all treatment conditions. Vertical reference lines denote median circularity values. **(F)** Fold change in organoid morphological complexity categories relative to DMSO control. Data are shown as mean ± SD. Statistical significance was assessed using unpaired Welch’s *t*-tests (*p* < 0.05, \**p* < 0.01).

Baseline organoid formation was observed from untreated and control vehicle-treated (DMSO) organoids that exhibited a unimodal organoid size distribution centered around medium (>12,000-33,000um^3^) to large (>33000 – 90000 µm^3^) structures that were highly complex comprised of branched TLDUs (Fig. 1e, Supplemental Figure 1c). In contrast, exposure to structurally diverse chemicals resulted in a broad spectrum of developmental abnormalities, ranging from impaired organogenesis to mild or no phenotypes (Fig. 1e,f; Supplemental Table 2). 2,4-DHBP, Kynurenine, PFOA, and S-equol all resulted in reductions in organoid formation, size distribution shifts toward smaller organoids, with a reduction in architectural complexity suggesting these exposures might be leading to more generalized developmental and morphogenic disruption.

The other chemicals produced discordant developmental phenotypes. Some reduced organoid formation without major changes in size or complexity; others altered size or complexity without a reduction in organoid numbers (Fig. 1c-f, Supplemental Table 2). This suggested that such chemicals are affecting different stages and biological processes during development. Genistein and BP-3 both produced strong architectural and size disruptions shifts but showed no effects on overall organoid number. In contrast, BP showed significant decrease in organoid formation but no effect on size or complexity, while BPP resulted in significant loss of organoid numbers, some effect on size, but little effect on complexity. Several other bisphenols (e.g., BPA, BPS, BPF, and BPAF), as well as benzylparaben, and PFOS exhibited both shifts in organoid size and in morphogenic complexity, but only modest changes in organoid numbers. Together, these findings indicate that organoid formation, size, and architectural complexity represent overlapping but non-redundant aspects of human mammary development that can be affected by chemical exposures. Furthermore, these results show that EDCs exert heterogeneous effects on breast organoid development that selectively disrupt processes impacting developmental yield, growth dynamics, or tissue organization.

### Physiologic Doses of Bisphenol Mixtures Disrupt Breast Organoid Development Toward ECM-Remodeling and EMT-Associated States

Humans are exposed to complex mixtures of environmental chemicals rather than single agents, with biomonitoring studies demonstrating the simultaneous presence of multiple EDCs and at lower doses^5,21^. These co-exposures may produce biological effects that are not predictable from single-agent studies at higher concentrations, highlighting the need to capture combinatorial effects at physiological concentrations. The observation that individual bisphenols induce distinct developmental phenotypes suggests that these compounds may disrupt different aspects of mammary organogenesis. Given their widespread detection, prevalence, and frequent co-occurrence in the general population^26^, we therefore evaluated a mixture composed of BPA, BPS and BPF to determine whether combined exposure reveals developmental effects not observed in single-agent assays. A 5 nM concentration for each chemical was selected to approximate environmentally relevant exposure levels reported in human biomonitoring studies^1,5^.

We treated 3D hydrogel cultures established from single cells obtained from reduction mammoplasty tissue (ages 17–31, Supplemental Table 3) from different patient donors (N=6) with combined bisphenols BPA, BPS, and BPF (3BPX) continually for 21 days and scored developmental and morphological disruption (Fig 2a, Supplemental Figure 2). Significant patient heterogeneity was observed in response to organoid formation and complexity in response to exposure. Nearly all donors (5/6) exhibited decreased organoid formation compared to vehicle-treated control gels. Precocious lobule formation was observed in two donor samples, and one donor sample resulted in the formation of abnormal organoids (Supplemental Fig. 2). Given the extensive patient heterogeneity responses, we established a semi-quantitative composite developmental disruption index (DDI) score to capture organoid developmental disruption more generally, incorporating the three phenotypic criteria metrics – organoid number, organoid complexity (% lobule formation), and abnormal organoid number – across all patient samples into a single score. When analyzing for developmental disruption, all donors demonstrated at least one phenotype and there was a statistically significant difference in DDI in response to continuous low dose 3BPX exposure (Fig. 2A).

**Figure 2.**
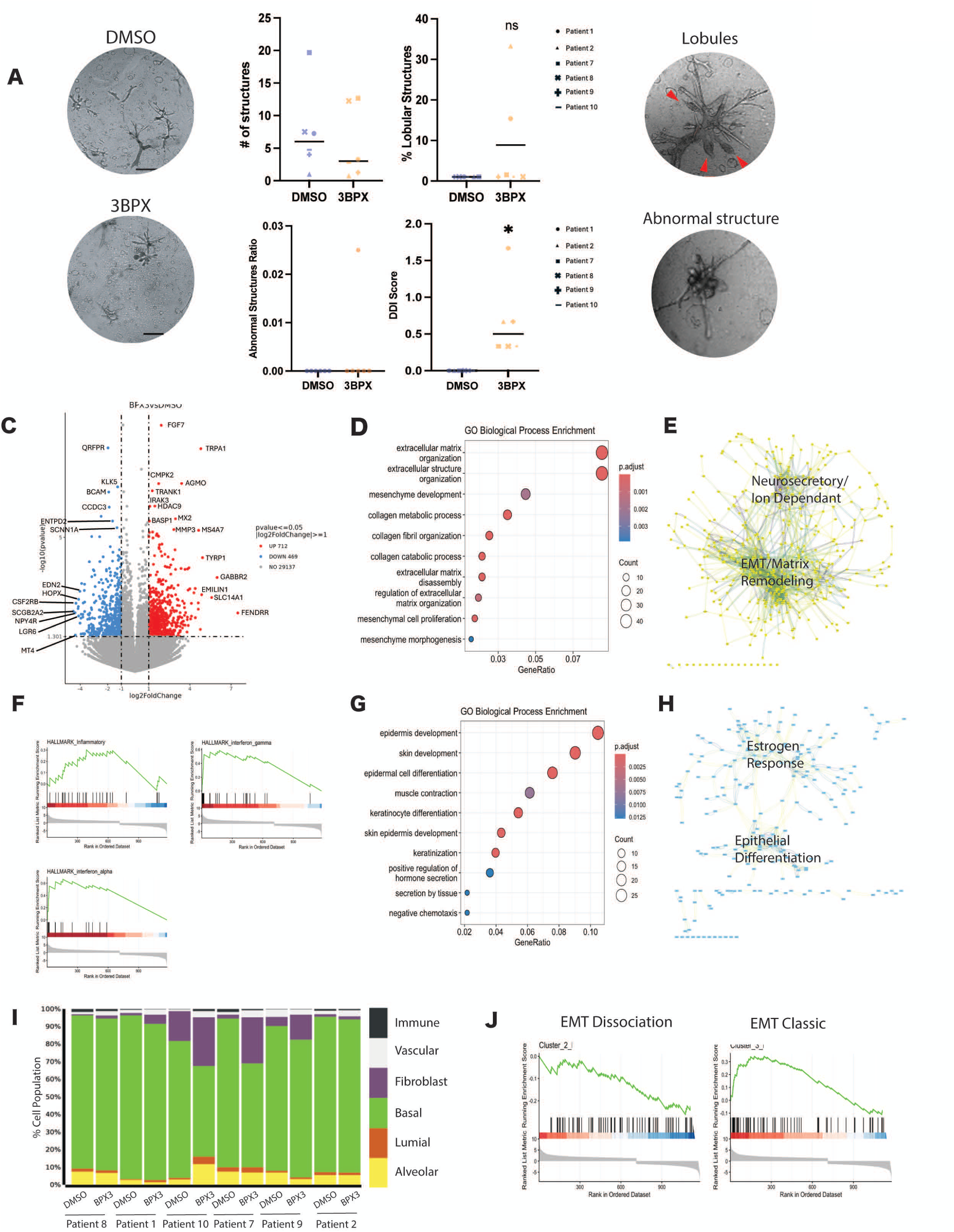
Bisphenol mixture (3BPX) exposure induces morphological disruption and transcriptional reprogramming in human breast organoids. **(A)** Representative bright-field images illustrating distinct organoid morphological phenotypes following exposure to DMSO or 3BPX (scale bar, 500 µm), with corresponding quantitative morphometric analyses of organoids cultured for 14 days from six independent patient donors. Statistical significance was assessed using the Wilcoxon signed-rank test. **(B)** Volcano plot of differentially expressed genes identified by bulk RNA sequencing of organoids exposed to 3BPX for 21 days compared with DMSO controls (n = 6 patient donors). Log₂ fold change is plotted on the x-axis and −log₁₀ *p*-value on the y-axis; significantly upregulated and downregulated genes are highlighted. **(C, D)** Gene Ontology (GO) biological process enrichment analysis of genes upregulated protein–protein interaction (PPI) network analysis of differentially expressed genes visualized using Cytoscape. Networks include high-confidence interactions (STRING score 400–700). Key hub genes and major functional modules are annotated. **(E)** Gene set enrichment analysis (GSEA) demonstrating enrichment of inflammatory–related transcriptional programs in 3BPX-treated organoids. **(F,G)** GO enrichment analysis and PPI network analysis of downregulated genes following 3BPX exposure (adjusted *p* < 0.05). Key hub genes and major functional modules are annotated. **(H)** Cell-state deconvolution of bulk RNA-sequencing data using CIBERSORTx, showing relative proportions of epithelial, stromal, and immune-associated cell states within organoids at day 21. Bar plots represent the percentage contribution of each cell population per sample. Quantification of the fibroblast/EMT-associated population is shown (right); statistical significance was assessed using the Wilcoxon test. **(F)** Gene set enrichment analysis (GSEA) demonstrating enrichment of epithelial–mesenchymal transition (EMT)–related transcriptional programs in 3BPX-treated organoids. Enrichment scores were calculated using the GSEA R package based on ranked log fold-change values.

Given this finding, we wanted to determine whether this disruption was primarily driven by any single bisphenol. Accordingly, hydrogels seeded with different donor-derived samples were exposed to 5 nM BPA, BPS, or BPF as single agents and monitored for organoid development. Exposure to any single bisphenol did not significantly alter organoid development or affect organoid number or complexity (Supplemental Figure 2) indicating that developmental phenotypes emerging from 3BPX exposure is due to the combined response of the mixed bisphenols rather than any single individual component.

We also examined the transcriptional changes due to 3BPX exposure by RNAseq from organoids treated continuously with 3BPX or vehicle control from the same 6 donors. Differentially expressed genes (DEGs) included 712 up-regulated genes and 469 down-regulated genes, with FGF7 and MMP3 among the most highly induced genes observed in response to 3BPX exposure across all 6 donor-derived organoids (Fig. 2B). Gene Ontology and pathway enrichment analyses revealed that up-regulated pathways were dominated by ECM remodeling processes, including matrix metalloproteinase activity and collagen metabolism, while down-regulated genes were enriched for epithelial differentiation, breast-specific markers, and cell-cell junction organization (Fig. 2C-H). Two dense clusters of highly connected hub genes were observed to infer higher-order protein network organization of functional and cellular relationships among up-regulated DEGs. A major hub involving ECM remodeling, epithelial-to-mesenchymal transition (EMT), and a second notable interaction cluster consisted of upregulated genes involved in neurosecretory-like vesicle and ion-dependent signaling (Fig. 2E, 2F, Supplemental Figure 3, Supplemental Table 4). Interestingly, within the EMT cluster, we observed gene set enrichment for inflammatory and interferon (IFNα/γ) responses (Fig. 2E, 2F, Supplemental Figure 3) suggestive of an EMT program that controls invasion and inflammation as observed in cancer ^45^. Among down-regulated transcripts, networks were observed related to estrogen-responsive epithelial differentiation and barrier-formation programs (Fig. 2H, Supplemental Table 4). Genes in this cluster included TGM1, CASP14, KRT10, KLK5, ERBB3, and DSC3, consistent with attenuation of epithelial differentiation programs.

Given the prominent EMT and matrix remodeling signature, we used lineage-deconvolution to infer whether any population-level changes in cell lineages might provide insights into whether the gene expression changes observed from 3BPX exposure might be due to shifts in the proportion of cell lineages within the organoids^27^. While there were no shifts in luminal or basal epithelial cell lineages, a significant change in the proportion of fibroblast/EMT cell populations was observed in organoids exposed to 3BPX (Fig. 2I)

The concurrent downregulation of epithelial differentiation, upregulation of ECM remodeling genes, and expansion of fibroblast/EMT cell lineages is consistent with the notion that 3BPX induces the EMT program. The EMT triggers several distinct biological processes including epithelial cell dissociation, cell plasticity, and invasion and migration. These different contexts can be transcriptionally deconvoluted into a spectrum of transcriptional programs: i) stemness genes coupled to cellular plasticity and self-renewal, ii) dissociative genes associated with loss of epithelial cohesion and tissue integrity, iii) classic signaling genes associated with canonical developmental pathways, iv) neural genes related to neural crest-like programs, and v) other EMT related genes that don’t fit canonical categories^28^. GSEA analysis was used to assess which of the EMT programs were induced by 3BPX exposures. Notably, the dissociative and classic EMT programs defined by the downregulation of junctional proteins, ECM remodeling and wound response signaling, and increased epithelial plasticity without full mesenchymal conversion were statistically enriched in the 3BPX signature (Fig. 2J, Supplemental Table 4). Taken together, the transcriptional changes suggest that continuous 3BPX exposure induces (i) reduction in breast epithelial identity and adhesion programs and (ii) activation of ECM remodeling pathways promoting epithelial plasticity and ECM-remodeling states during organoid development.

### Mixed Bisphenol-Exposure Signature is enriched in Human Invasive Lobular Breast Cancer and Maps to a Unique Cancer Cell State

To determine whether the 3BPX signature exists in human breast cancers, we projected the bulk RNA-seq datasets from TCGA-BRCA, METABRIC, and SCAN-B onto the 3BPX signature. Across cohorts, BPX3 scores varied by intrinsic subtype, with higher scores in Luminal A and Normal-like tumors and lower scores in Basal-like tumors, indicating that the BPX3 transcriptional program is associated with Luminal tumors. Notably, BPX3 scores were significantly higher in a histological subtype of Luminal tumors, invasive lobular carcinoma (ILC), compared to invasive ductal carcinoma (IDC). Within Luminal A tumors, BPX3 scores were consistently higher in ILC across cohorts, with a modest but reproducible effect size. In SCAN-B, the relationship varied across subtypes, with higher BPX3 scores in ILC within Luminal A and Normal-like tumors, but attenuated or absent differences in Luminal B and HER2-enriched tumors. Because ILC is predominantly estrogen receptor–positive (ER⁺), we evaluated whether this association was independent of subtype composition. The ILC–IDC difference persisted within ER⁺ tumors and remained significant after adjusting for PAM50 subtype, with similar results upon inclusion of ER status as a covariate, indicating that BPX3 enrichment in ILC is not solely explained by ER status or subtype distribution.

ER^+^, ILC is characterized by loss of E-cadherin, and the adoption of an invasive, partial EMT epithelial state ^31^. Furthermore, tumor cellularity is typically low in ILC, with the stroma often comprising the majority of the tumor mass due to its diffuse infiltrative growth pattern and reduced inflammatory infiltrates ^32^. Given that ILC tumors are frequently stroma-rich and tumor cell-poor we next examined whether the 3BPX-derived signature might map onto single-cell RNA-seq tumor profiles. We leveraged a large human breast cancer atlas^33^ (>600,000 cells; 138 patients; 8 datasets) spanning tumors annotated across PAM50 subtypes to enable highly resolved characterization of epithelial, immune, and stromal heterogeneity defined across the PAM50 (Luminal A, Luminal B, Basal, Normal, and Her2) breast cancer subtypes (Fig. 3D,E).

**Figure 3.**
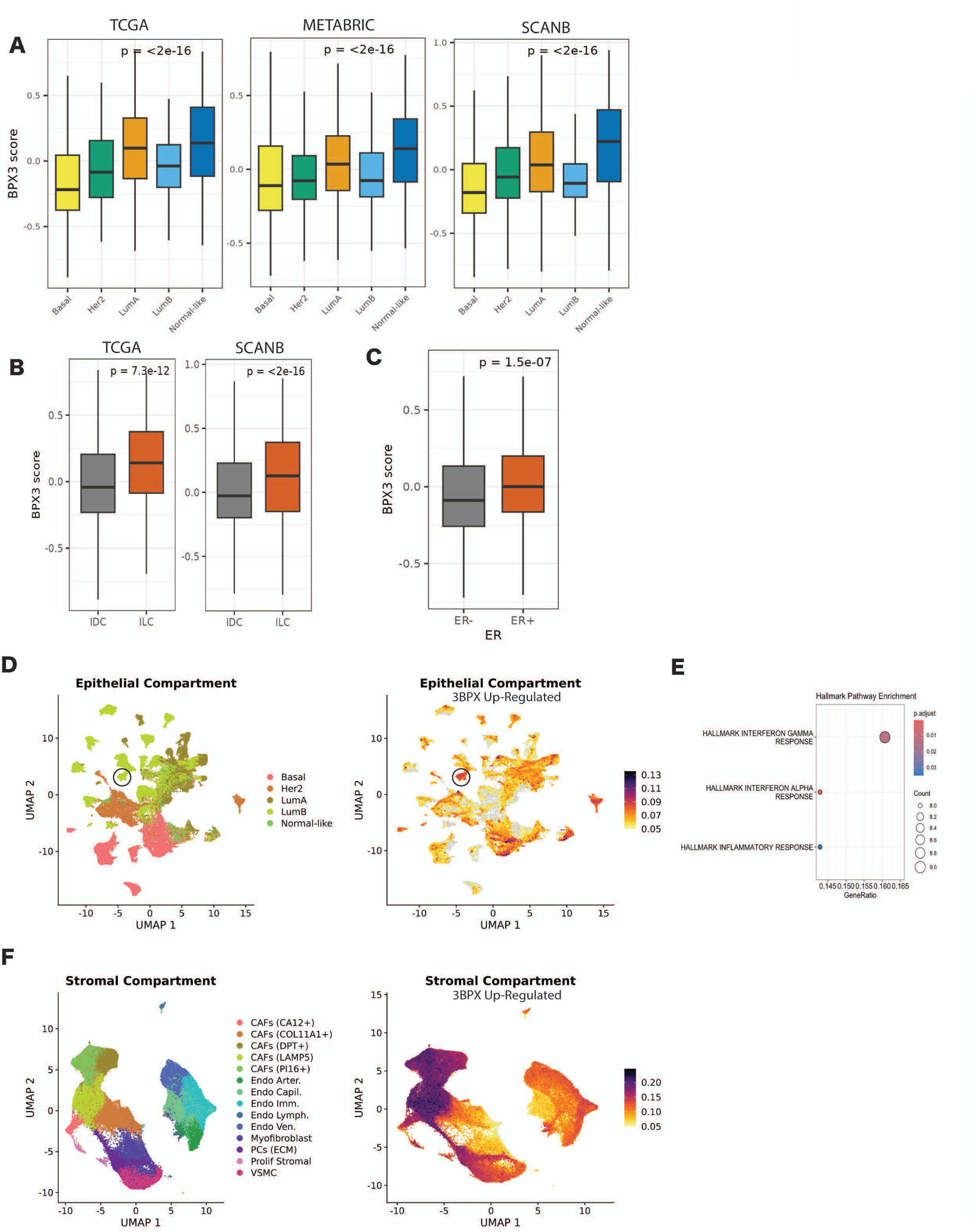
Mixed bisphenol exposure signature enrichment across breast cancer subtypes and mapping to defined epithelial and stromal cell states. **(A)** Distribution of the 3BPX gene expression signature score across intrinsic breast cancer subtypes in three independent cohorts: TCGA, METABRIC, and SCAN-B. Tumors are stratified by PAM50 subtype (Basal, HER2-enriched, Luminal A, Luminal B, and Normal-like). **(B)** Comparison of 3BPX signature scores between invasive ductal carcinoma (IDC) and invasive lobular carcinoma (ILC) in TCGA and SCAN-B cohorts. **(C)** Stratification of tumors by estrogen receptor (ER) status. **(D)** Single-cell RNA-sequencing (scRNA-seq) atlas of human breast cancers showing the epithelial compartment annotated by intrinsic cancer subtype (left). Genes upregulated following 3BPX exposure in breast organoids (bulk RNA-sequencing; n = 6 donors) were projected onto the epithelial atlas to identify cancer cell states exhibiting concordant transcriptional programs (right). Color intensity indicates the degree of enrichment of the 3BPX transcriptional signature within each epithelial cell population. Circle denotes cluster region of highest enrichment for the 3BPX transcriptional signature. **(E)** Gene set enrichment analysis (GSEA) of Hallmark pathways significantly enriched (*p* < 0.05) within the Luminal B epithelial subpopulation most strongly associated with the 3BPX transcriptional signature. Dot size reflects the number of overlapping genes between the Luminal B subpopulation and each Hallmark pathway, and color indicates statistical significance. **(F)** scRNA-seq atlas of human breast cancers showing the stromal compartment annotated by stromal cell type (left). Upregulated genes from 3BPX-treated organoids were similarly projected onto the stromal atlas to identify stromal cell populations exhibiting enrichment of the 3BPX transcriptional signature (right).

Notably, expression of the BPX3 signature, as quantified by AUCell^46^, localized to a discrete subset of epithelial cells within Luminal B cell state (Fig. 3D), rather than being broadly distributed across all luminal cancers or across all Luminal A or Luminal B epithelial states. Although we did not identify a single pathway or category that uniquely separated this cluster from other epithelial populations, genes contributing to this cell state that mapped to BPX3 exposure were significantly enriched for interferon (IFNα/γ) response programs (Fig. 3E). The ER^+^ subtype-level tumor cell state localization complements the bulk analyses and is consistent with a BPX3 exposure signature that is enriched for a stress and immune-responsive epithelial state found within an ER^+^intrinsic subtype rather than uniform expression.

Given the strong EMT signature due to 3BPX expression, we also examined whether it maps to stromal compartments within the same cancer atlas. Tumor stroma largely resolves into fibroblast and endothelial cells (e.g., VMSC, vascular, lymphatic) (Fig. 3F, Supplemental Table 5) ^33^. Fibroblasts are further resolved into six clusters corresponding to previously defined cancer-associated fibroblast (CAF) states and myofibroblasts (Fig. 3F) ^33^. Projection of the stromal cells’ profiles onto the 3BPX signature reveals selective enrichment mainly within PI16⁺ fibroblasts and LAMP5⁺ CAF subsets, rather than across all stromal or fibroblast populations. Notably, PI16⁺ fibroblasts represent a population closely related to normal tissue fibroblasts ^34^, while LAMP5⁺ CAFs, are characterized by expression of LAMP5 and cystatins, and implicated in promoting EMT ^35–37^. Given these findings, we next examined whether the BPX3 signature might be enriched in the stromal compartment from laser-capture microdissected ILC ^38^. Indeed, ∼18% (8/45) of the genes unique to ILC stroma overlapped with BPX3 genes (ADAMTS18/EFEMP2/MMP14/PAPPA/TIMP2/MMP2/ITGA10/COL6A3) suggesting, BPX3 exposure program recapitulates core stromal features of human ILC. Collectively, these data suggest the transcriptional changes due to BPX3 exposure are enriched in ILCs, maps to both discrete stress and immune-responsive epithelial states and specific stromal and EMT states in breast cancer. This compartmentalization suggests that environmental exposures may contribute to tumorigenesis not only through epithelial-intrinsic effects, but also through remodeling of the tumor microenvironment.

### Exposure to Complex Chemical Mixtures Results in Developmental Disruption

The observation that exposure to 3BPX induced transcriptional changes associated with a dissociative EMT program was reminiscent of findings that showed benzo[a]pyrene (BaP) exposure, a well-characterized environmental carcinogen, also elicits epithelial barrier disruption and EMT in a 3D human bronchial epithelial model^29^. We therefore examined whether the transcriptional program induced by 3BPX exposure might overlap with those induced by exposure with BaP. Indeed, 70 genes were shared between the 3BPX signature and the BaP signature (Fig. 4A) despite coming from independent epithelial systems and different chemical exposures. These overlapping genes were enriched for pathways related to ECM remodeling and EMT.

**Figure 4.**
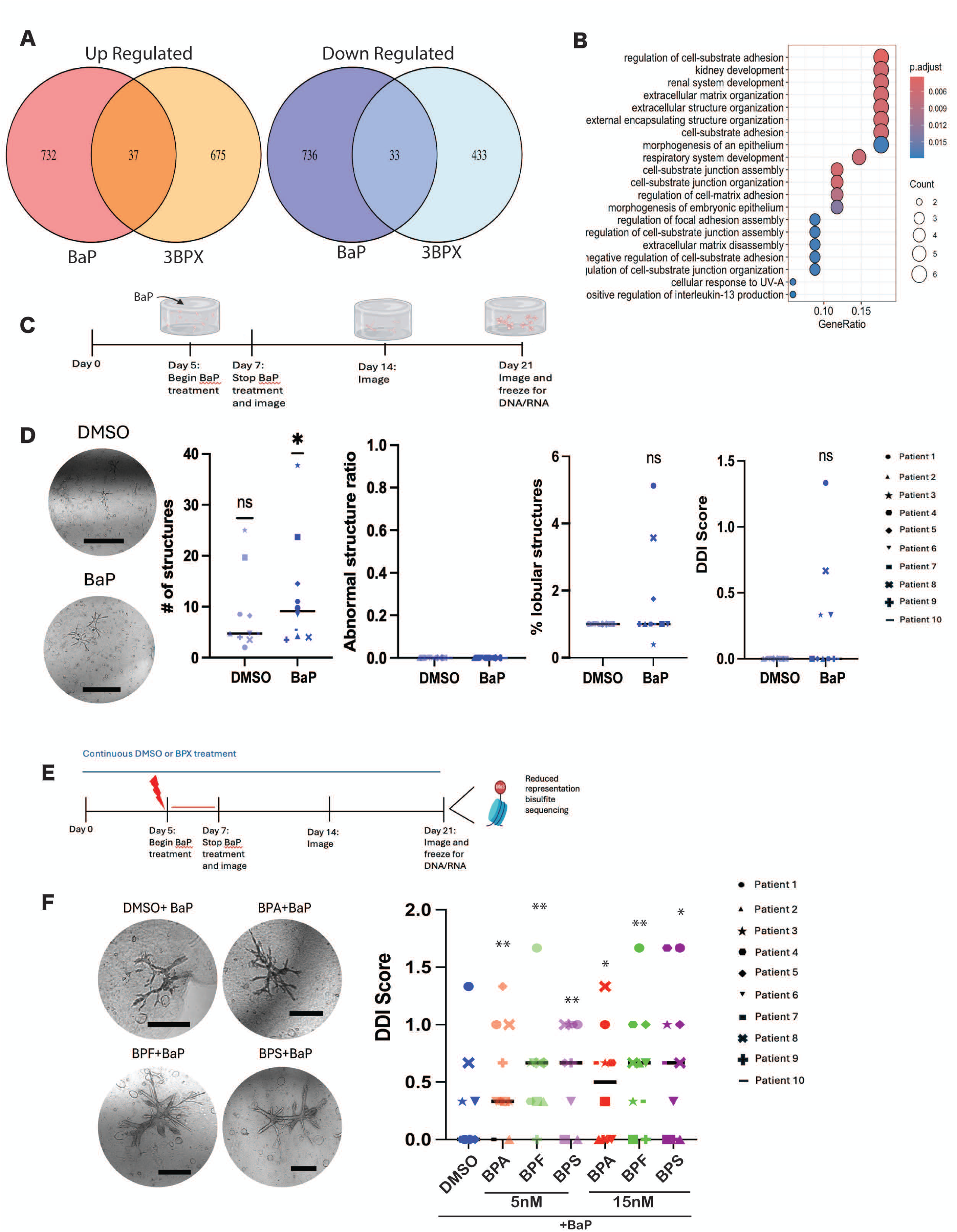
Distinct environmental exposures show convergent molecular and developmental effects. **(A)** Venn diagrams showing overlap of significantly upregulated (left) and downregulated (right) genes identified by bulk RNA sequencing following exposure to benzo[a]pyrene (BaP) or a mixed bisphenol formulation (3BPX). BaP data were generated from primary human bronchial epithelial organoids, while 3BPX data were generated from primary human breast epithelial organoids. Statistical significance of gene set overlap was assessed using Fisher’s exact test. **(B)** GO analysis demonstrating enrichment of overlapping dissociative EMT–related transcriptional programs between 3BPX-treated organoids and BaP-treated bronchial epithelium in 3D.**(C)** Schematic illustrating the experimental timeline for BaP exposure during organoid development, including treatment initiation, imaging, and endpoint collection for downstream DNA and RNA analyses. **(D)** Representative bright-field images of organoid morphology following BaP exposure (scale bar, 500 µm), with corresponding quantitative morphometric analyses of organoids cultured to day 14 from ten independent patient donors. Statistical significance was assessed using the Wilcoxon signed-rank test. **(E)** Schematic illustrating the experimental timeline for organoid exposure to individual bisphenols (BPA, BPF, or BPS) in the presence or absence of BaP, including treatment initiation, imaging, and endpoint collection for DNA and RNA analyses. **(F)** Representative bright-field microscopy images of organoids for each treatment condition (left; scale bar, 500 µm) with corresponding semi-quantitative developmental disruption index (DDI) scores (right) from ten independent patient donors. Each symbol represents an individual donor.

Given this, we next assessed whether exposure to BaP might also lead to similar developmental alterations during organoid formation as 3BPX. Human breast epithelial cells were seeded into 3D hydrogels and treated on day 5 with BaP and assessed by immunofluorescence for γH2AX and Zombie green to assess DNA damage and viability. Day 5 corresponds to a developmental transition phase characterized by active patterning and expansion of progenitor populations that give rise to ductal and lobular structures^24,25^ (Supplemental Fig. 1A). Exposure during this window enables perturbations to propagate through subsequent developmental lineages^25^. The dose of 1uM was established as it induced robust DNA damage by 48 hours but limited cell death (Supplemental Fig. 3B).

Single human breast epithelial cells were seeded into 3D hydrogels from 10 different patient donors and treated on day 5 with 1uM BaP and allowed to develop into organoids over the next 17 days in the absence of BaP. Similar to 3BPX exposure, single dose exposure to BaP demonstrated extensive heterogeneity across 10 different donors, with increased organoid formation and precocious lobule formation (Fig. 4D, Supplemental Figure 3). While both BaP and 3BPX exhibited similar changes in lobule complexity, 3BPX differed from BaP in both organoid formation and in the formation of abnormal organoids (Fig. 2). BaP increased organoid formation but did not cause formation of abnormal organoids, whereas 3BPX reduced organoid formation and did produce abnormal organoids. Together these results indicate that BaP and 3BPX, with different primary modes of action, exhibit partial overlapping developmental phenotypes and transcriptional programs.

Humans are exposed to simultaneous complex mixtures of environmental chemicals rather than mixtures of agents within the same chemical class ^21,30^. These complex co-exposures may produce combinatorial biological effects that are not predictable from single class studies. We subsequently examined the effect of continuous low dose single agent bisphenols combined with transient co-exposure of BaP on organoid development. Primary human breast epithelial cells derived from independent donors (N=10) were seeded into 3D hydrogels and exposed continuously to individual bisphenols (5 or 15 nM) throughout organoid development. BaP was introduced transiently during days 5–7, corresponding to a developmental transition phase characterized by active patterning and expansion of epithelial progenitor populations, after which it was withdrawn while bisphenol exposure was maintained for the remainder of the culture period (Fig. 4E). This design enabled assessment of continuous non-genotoxic exposures interacting with a temporally restricted DNA-damaging insult during development.

Consistent with prior observations, substantial inter-donor variability was observed across all treatment conditions (Fig. 4D). However, combined exposures were associated with increased organoid formation, increased frequency of precocious lobule development, and the emergence of abnormal organoid structures characterized by disorganized or clustered lobules with DDI score significantly increased following co-exposure to bisphenols compared to BaP alone (Fig. 4F). These findings suggest that low-dose bisphenol exposure can modulate the developmental response to an additional environmental insult, highlighting the importance of mixture effects in shaping developmental outcomes.

### Chemical Exposures Induce Unique DNA Methylation Fingerprints

The results above indicate that exposure to environmental compounds during organoid development can induce durable changes in tissue architecture. These developmental changes could prime a tissue to the formation of disease states or pre-neoplastic lesions. Since the molecular basis of cancer susceptibility to chemical exposure remains poorly understood, and cannot be fully explained by genetic mutations or a single etiologic pathway, we examined whether epigenetic changes associated with chemical exposure in developing organoids might explain susceptibility to malignant transformation. Accordingly, we performed whole-genome DNA methylation profiling to examine these different environmental exposures on the epigenome. Whole genome DNA methylation analysis was performed using reduced representation bisulfite sequencing (RRBS) on pooled DNA collected from 4 gels per donor across all 10 patient donors (N=40 samples per treatment) on 21-day organoids that had been treated continuously with 3BPX or single bisphenols with a transient 48hr exposure with BaP. Pooling the DNA from each chemical exposure across all donors reduces underlying patient heterogeneity to maximize the detection of methylation differences that would be specific to the chemical exposure and not the individual donor sample.

Genome-wide analysis revealed extensive differentially methylated regions (DMRs) following both BaP and 3BPX treatment relative to vehicle-treated controls distributed across all chromosomes (Fig. 5A-D). While both exposures produced thousands of DMRs, overlap between BaP- and 3BPX-associated sites was limited, indicating that these agents induce largely distinct epigenetic programs rather than a shared, nonspecific methylation response. DMRs were localized to both gene bodies and distal regulatory regions as well as promoter-proximal elements (Fig. 5E, Supplemental Figure 3). At individual loci, both hypermethylation and hypomethylation were observed, with directionality varying by gene rather than following a uniform exposure-wide pattern.

**Figure 5.**
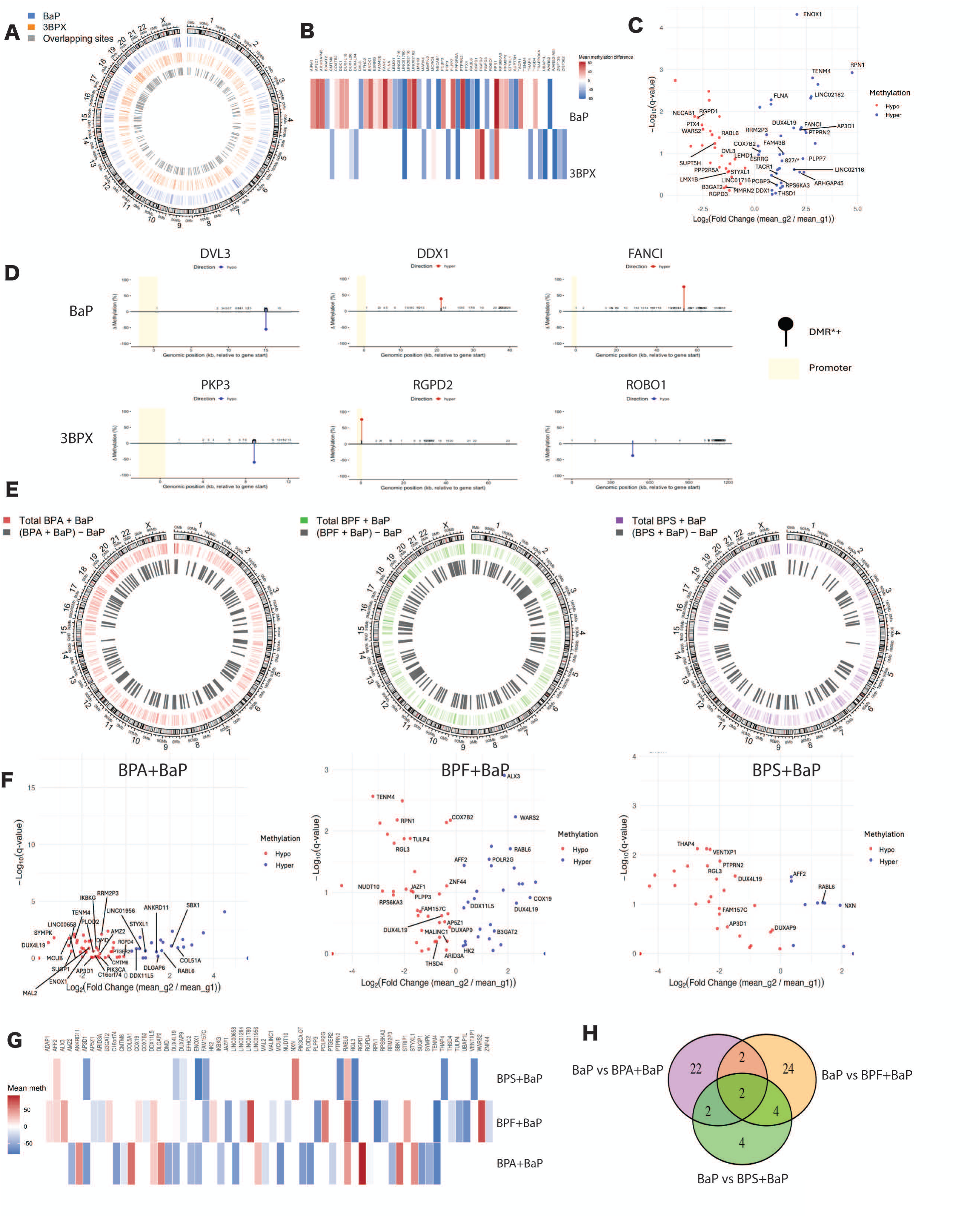
Exposure to complex chemical mixtures amplifies developmental disruption and induces distinct epigenetic reprogramming. **(D)** Circos plot summarizing differentially methylated regions (DMRs) identified by reduced representation bisulfite sequencing (RRBS) following exposure to BaP (outer ring, blue) or 3BPX (middle ring, orange), each compared to DMSO controls and mapped across the genome by chromosome. The inner gray ring denotes DMRs shared between both exposure conditions. **(E)** Lollipop plots depicting the genomic localization of high-confidence DMRs mapped to annotated gene features. Only DMRs within the top 10% of ranked regions based on significance scores were included. Lollipop height reflects the magnitude of methylation change relative to DMSO control. Blue lollipops indicate hypomethylation and red lollipops indicate hypermethylation. **(F)** Volcano plot of DMRs identified from RRBS analysis of pooled DNA from primary organoids exposed to BaP and harvested at day 21 from ten patient donors. Log₂ fold change in methylation is plotted on the x-axis and −log₁₀ *q*-value on the y-axis. **(C)** Circos plots summarizing differentially methylated regions (DMRs) identified by reduced representation bisulfite sequencing (RRBS) following co-exposure to BaP with BPA (red), BPF (green), or BPS (purple). Analyses were performed on pooled DNA from ten patient donors per condition. Inner gray rings denote DMRs unique to each bisphenol + BaP condition after subtracting regions shared with BaP-only exposure. **(D)** Volcano plots of DMRs mapping to annotated genes for each bisphenol + BaP condition. Log₂ fold change in methylation relative to DMSO controls is shown on the x-axis and −log₁₀ *q*-value on the y-axis. Points are colored by direction of methylation change (hypermethylated or hypomethylated). **(E)** Venn diagram depicting overlap of gene-associated DMRs identified across BPA + BaP, BPF + BaP, and BPS + BaP exposure conditions.

Mapping DMRs to gene regions revealed methylation changes affecting genes involved in epithelial organization, cell polarity, immune signaling, and chromatin-associated regulation of cell state, including *PKP3, MARK4, MORC4, RPN1*, *CMTM6*, and multiple *DUX4* family members for 3BPX and DVL3, *RPS6KA3, SUPT5H, DDX*, and *FANCI* for BaP (Fig. 5C-D). A subset of loci, including *GPD1, LINC01780, PPP2R5A,* and *RPN1*, exhibited DMRs in both 3BPX- and BaP-treated samples, indicating partial overlap between exposure-associated methylation changes. Notably, methylation changes often spanned extended genomic regions rather than being restricted to discrete promoter sites (Fig. 5D, Supplemental Fig. 3), consistent with broad epigenetic remodeling. Collectively these observations raise the possibility that distinct environmental exposures may converge on shared developmental and transcriptional programs while establishing exposure-specific epigenetic changes.

Genome-wide distributions of DMRs on complex chemical exposures also revealed each bisphenol–BaP condition exhibited widespread methylation changes distributed across all chromosomes. While a substantial number of DMRs were observed across all conditions, the genomic distribution and density of DMRs varied between bisphenol–BaP combinations, suggesting that each exposure condition is associated with a distinct epigenetic profile, consistent with context-dependent differences in epigenetic remodeling (Fig. 5E-H). There was a broad distribution of both hypermethylated and hypomethylated regions across all conditions, with statistically significant changes detected despite relatively modest effect sizes. While a subset of gene-associated DMRs was shared between conditions, a substantial proportion of DMRs were unique to each bisphenol–BaP combination (Fig 5E, F). While each bisphenol–BaP condition retained a distinct set of unique DMRs, *RABL6* and *AP3D1* gene-coding DMRs were identified across all conditions, including single BaP exposure (Fig. 5H) defining a gene region specific methylation signature to BaP exposure. Together these findings suggest that while BaP drives a shared component of the epigenetic response, bisphenols modulate this response in a compound-specific manner. Furthermore, these findings indicate that individual bisphenols do not produce interchangeable epigenetic outcomes when combined with BaP, but instead generate distinct gene-associated methylation signatures. Together, these results suggest that combined environmental exposures produce both shared and exposure-specific epigenetic programs, resulting in heterogeneous but reproducible patterns of DNA methylation across the genome.

### Human Breast Cancers Retain a DNA Methylation ‘Scar’ of EDC Exposure

We next evaluated whether these chemical-induced DNA methylation fingerprints during breast organoid development are detectable in human breast tissue. We projected RRBS-derived methylation signatures onto TCGA-BRCA methylation data comprising 784 primary tumors and 97 adjacent normal breast samples. Notably, Normal-like and Basal tumors exhibited significantly higher enrichment of exposure-derived methylation signatures compared to other cancer subtypes, with Normal-like breast cancers enriched for 3BPX exposure while Basal tumors enriched for complex BaP+BPS exposure (Fig 6A, Supplemental Figure 5A). These findings indicate that exposure-induced DNA methylation fingerprints are differentially represented across breast cancer subtypes, suggesting that environmental exposures may contribute to shaping subtype-specific epigenetic states.

**Figure 6.**
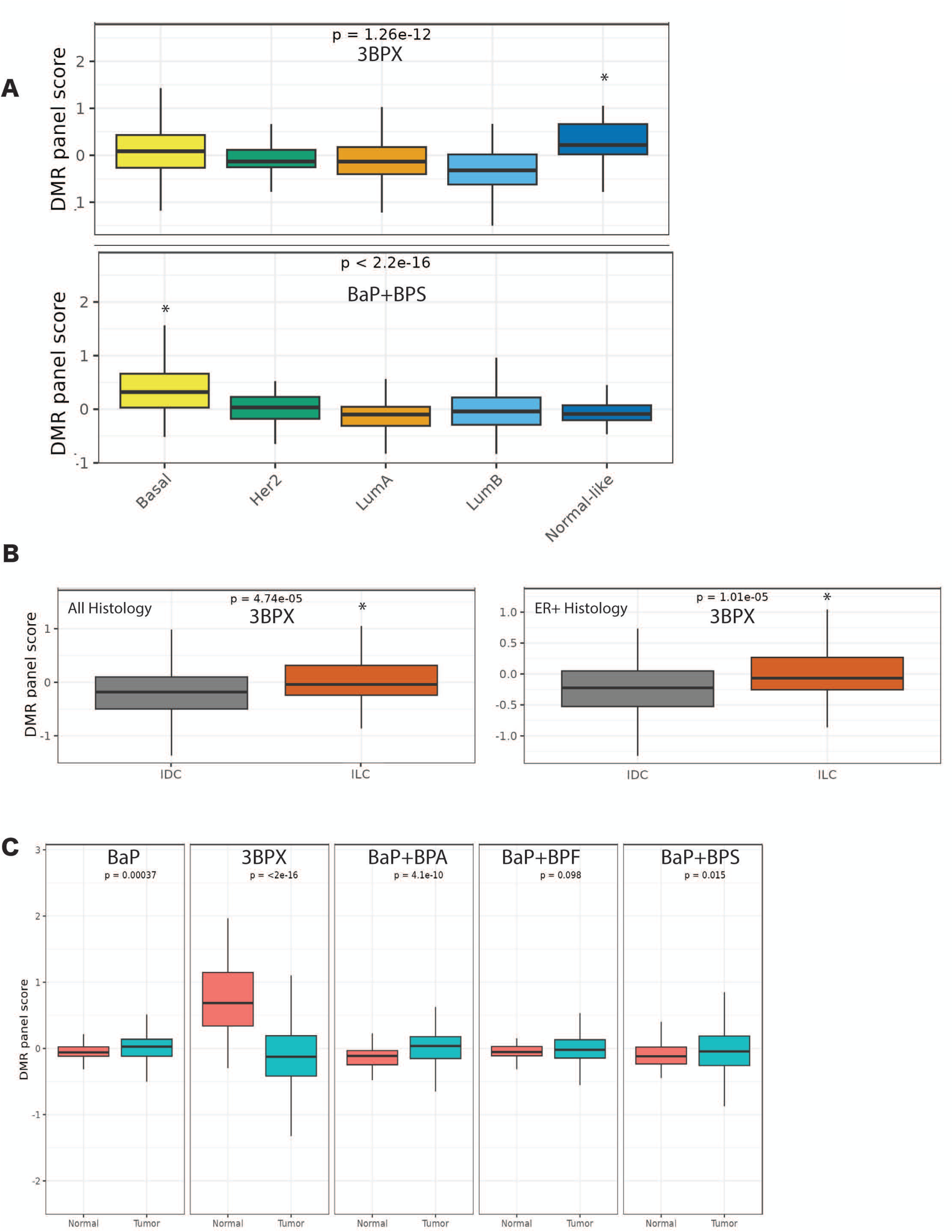
Exposure-derived DNA methylation signatures from organoids are detectable in human breast cancer. **(A)** Projection of RRBS-derived methylation signatures onto TCGA-BRCA DNA methylation data. The 3BPX-derived methylation signature is enriched in Normal-like tumors, while the BaP+BPS-associated signature shows strongest enrichment in Basal tumors. Panel scores represent direction-aligned, z-scored methylation values aggregated across gene-associated DMRs measurable in TCGA (±500 bp overlap). **(B)** Association of exposure-derived methylation signatures with tumor histology. The 3BPX methylation signature is significantly elevated in invasive lobular carcinoma (ILC) compared to invasive ductal carcinoma (IDC), and this relationship persists when analysis is restricted to ER⁺ tumors. **(C)** Tumor versus adjacent normal comparison of exposure-derived methylation signatures. The 3BPX-derived methylation signature demonstrates the strongest and most consistent separation between tumor and adjacent normal tissue, with significantly lower scores in tumors. In contrast, other exposure-derived signatures (including BaP alone and bisphenol–BaP co-exposures) show weaker, inconsistent, or non-significant tumor–normal differences.

Given the strong association between ILC and the BPX3 transcriptional signature, we next examined whether the 3BPX-derived methylation signature was similarly associated with tumor histology. Indeed, 3BPX methylation signature scores were significantly higher in ILC compared to IDC, indicating that the 3BPX-associated epigenetic program is preferentially retained in lobular tumors (Fig. 6B, Supplemental Figure 5B,C). This difference persisted when analysis was restricted to ER⁺ tumors, demonstrating that the enrichment of the 3BPX methylation signature in ILC is not solely attributable to differences in ER status or subtype composition. Notably, no other exposure-derived methylation signatures retained significance after adjustment, identifying the 3BPX-derived methylation program as the most robust and independent epigenetic feature distinguishing lobular from ductal breast cancer.

Finally, we examined whether exposure-associated methylation signatures could distinguish tumor from adjacent normal tissue. The 3BPX-derived methylation signature demonstrated the most robust and consistent separation between tumors and adjacent normal samples (Fig. 6C), with a strong and coordinated shift in methylation across loci defining this panel. In contrast, methylation signatures derived from other exposure conditions, including BaP alone or bisphenol–BaP co-exposures, exhibited weaker or inconsistent tumor–normal differences, and in some cases there was no significant separation. These results suggest that the 3BPX-associated methylation program defines a stable epigenetic state that is detectable in normal tissue and systematically altered during tumorigenesis.

Together, these findings demonstrate that the 3BPX-exposure methylation signature is detected in human breast cancer and is enriched in Normal-like tumors and in ILC. The strong tumor–normal separation observed for this signature supports the presence of a persistent exposure-associated epigenetic “scar” linking developmental perturbation in organoid models to molecular features of human breast cancer. In contrast, single-agent and co-exposure conditions do not produce methylation programs that are equivalently represented across tumors.

## Discussion

Environmental exposures during critical windows of development are increasingly recognized as key determinants of lifetime cancer risk, yet the mechanisms by which EDCs affect the developing human breast remain poorly resolved ^9^. Our results show that chronic, low-dose exposures to chemical mixtures have the potential to alter cell state, disrupt tissue architecture, and induce persistent epigenetic changes creating an environment that may be more susceptible to malignant transformation. Current mutagenicity assays (eg. Ames test ^39^) restricts carcinogenic risk assessment to genotoxic mechanisms, overlooking the non-genotoxic pathways that have equally important effects on tissue development, cellular differentiation, hormonal signaling, or epigenetic programming. These latter processes are increasingly recognized as critical determinants of long-term cancer risk^40^.

A major advance of this study is establishing a 3D human breast organoid high-content screening (3D-HCS) platform to provide a human-relevant approach to assess non-genotoxic or genotoxic chemical exposure by directly measuring disruptions in development, tissue architecture, and cell fate programs linked to carcinogenic and developmental risk. Another major strength of this particular 3D hydrogel organoid model is its ability to capture multicellular tissue responses to environmental exposures due in part to the emergence of a stromal compartment that includes fibroblast-like and mesenchymal cell states in this model^24^. Notably, this stromal compartment was responsive to chemical exposure, revealing transcriptional and phenotypic programs associated with tumor CAFs, tissue remodeling, epithelial–stromal crosstalk, and injury responses that are increasingly recognized as central to non-genotoxic carcinogenic and developmental risk. The inclusion of stromal responses therefore enables detection of chemical effects that would be missed in epithelial-only systems. Together, this 3D hydrogel screening advancement could establish a standardized assay that could be widely adopted within regulatory toxicology frameworks to evaluate how chemical exposures during developmental windows reprogram tissues in ways that elevate cancer susceptibility later in life. Indeed, our findings show that continuous low-dose exposure to widespread EDCs, particularly bisphenol mixtures, induces reproducible morphological disruption, transcriptional rewiring, and durable epigenetic reprogramming during human mammary morphogenesis. These findings provide direct experimental evidence supporting a developmental origins model of breast cancer susceptibility^12^ in which chemical exposures create persistent epithelial vulnerabilities long before tumor initiation.

Our results also demonstrate that chemically distinct exposures produce distinct developmental phenotypes, yet converge on shared transcriptional programs characterized by wound response, inflammation, and EMT. This convergence suggests that diverse environmental insults engage a common stress-response axis that may represent a central node of vulnerability. Notably, mixture exposures elicited more severe phenotypes, highlighting that combined exposures, rather than single agents, may pose greater biological risk in real-world settings.

The finding that a mixed bisphenol exposure signature was enriched in ER+ Luminal tumors and especially in ILC was particularly notable given that the incidence rates for ER+ Luminal cancers have increased more steadily compared to other breast cancers across all racial and ethnic groups^42.^ Moreover, the incidence of ILC in particular has more than doubled since 1975, rising three times faster than other breast cancers ^43^ and climbing at the same rate in younger women (<50) as in older ones. This suggests a purely age-dependent accumulation of risk and is consistent with our findings that continuous exposures to EDCs leads to gene expression programs of ECM remodeling, dissociative EMT, and epithelial plasticity that are characteristics of ILC. The integration of morphologic disruption, EMT activation, and persistent epigenetic remodeling is consistent with a field cancerization model, in which histologically normal tissue acquires molecular alterations that increase susceptibility to transformation. Chronic EDC exposure might act as modifiers of epithelial cell state through gradual reprogramming of hormone-responsive epithelial cells toward adhesion-low, plastic states and primed for transformation.

At the epigenetic level, bisphenol mixtures established reproducible methylation changes across donors, affecting genes involved in nuclear organization, epithelial structure, and immune regulation. Notably, these exposure-associated marks were not confined to the experimental system, but were detectable in human breast tissue, where the 3BPX-derived signature exhibited strong separation between tumors and adjacent normal samples and was differentially represented across tumor subtypes. In particular, the 3BPX methylation program was enriched in Normal-like tumors and in invasive lobular carcinoma (ILC), where it remained associated with histology independent of ER status and intrinsic subtype, suggesting that this exposure-linked epigenetic state is selectively retained in specific tumor contexts. The observation that adjacent normal tissue exhibits higher similarity to the 3BPX-associated methylation program than tumors further indicates that tumorigenesis involves systematic remodeling of a pre-existing epigenetic state rather than the emergence of entirely de novo methylation patterns.

Together, these findings support a model in which environmental exposures coordinately reprogram epithelial tissues into a state characterized by altered architecture, increased plasticity, and modified immune interactions. Such changes may precede overt disease and establish a biologically primed field that is more susceptible to subsequent oncogenic events. In this framework, exposure-induced epigenetic programs are not uniformly retained across cancers, but instead align with specific tumor cell states and lineages, raising the possibility that environmental exposures may influence not only cancer risk, but also the phenotypic trajectory of tumor development. These observations are consistent with emerging evidence that epigenomic instability, rather than overt mutagenesis, may constitute an early event in environmentally promoted carcinogenesis^9,44^.

## Limitations

The use of primary human breast organoids enabled tissue-relevant mechanistic insight that circumvents the limitations of rodent models. However, a limitation of this approach is the heterogeneity observed at the level of patient heterogeneity but also at the individual organoid level, including differences in baseline morphology and growth kinetics. Thus, exposure-associated phenotypes may not generalize uniformly across all organoids or all donors. This intrinsic biological variability motivated the development of the Developmental Disruption Index (DDI), a quantitative framework designed to integrate diverse morphologic features into a standardized metric that enables systematic comparison of heterogeneous phenotypic outcomes. While follow-up validation in additional donors demonstrated both shared and donor-specific responses, expanding future screens across larger and more diverse donor cohorts will be important for refining exposure signatures and fully capturing population-level variability in susceptibility. Future work should expand the spectrum of chemicals and real-world mixtures tested. Despite these limitations, these findings provide a framework linking environmental exposures to altered human breast development, durable epigenetic reprogramming, and heightened susceptibility to transformation. By combining human organoid biology with AI-assisted screening, this work establishes a scalable platform for decoding how real-world chemical mixtures shape cancer risk.

## Methods

### Human breast tissue procurement and processing

Disease-free human breast tissue was obtained from discarded tissue of 13 different patients undergoing elective reduction mammoplasty or mastectomy procedures at Tufts Medical Center under protocols approved by the Tufts University Institutional Review Board (IRB #13521, Supplemental Table 3). All donors provided written informed consent prior to participation. Samples were de-identified in compliance with HIPAA regulations before transfer to the laboratory.

Tissue processing was performed as previously described^24–26^. Briefly, tissue was partially digested using 1.5 mg/mL collagenase (Millipore Sigma, 10103578001) and 18 µg/mL hyaluronidase (Sigma-Aldrich, H3506) to generate epithelial fragments. Fragments were washed and cryopreserved in 1 mL aliquots at −160 °C. For experiments, epithelial fragments were thawed and further dissociated into single cells using 0.25% trypsin (Thermo Fisher Scientific, 25200056) and 5 U/mL dispase (Roche, 4942078001).

### 3D hydrogel organoid culture

Single cells were embedded in 3D hydrogels as previously described^25,26^. Cells were seeded at 1,500 cells per 200 µL gel composed of 3.56 mg/mL collagen I (Millipore Sigma, 08-115), 10 µg/mL hyaluronic acid (Millipore Sigma, 385908), 40 µg/mL laminin (Gibco, 23017-015), and 20 µg/mL fibronectin (Sigma-Aldrich, F1056). Gels were plated into four-well chamber slides (Falcon, 354104) and polymerized for 1 h at 37 °C.

Phenol red–free mammary epithelial basal medium (MEBM; Lonza, CC-3152) was added and gels were gently released to float in media. Media were supplemented with bovine pituitary extract (52 µg/mL; Thermo Fisher Scientific, 13028014), hEGF (10 ng/mL; Sigma-Aldrich, E9644), insulin (5 µg/mL; Sigma-Aldrich, I9278), hydrocortisone (500 ng/mL; Sigma-Aldrich, H0888), GlutaMAX (1% v/v; Thermo Fisher Scientific), and antibiotic–antimycotic (1% v/v; Corning, 30-004-CL). Media were replaced every 2–3 days.

### Chemical exposures

Bisphenol A (BPA), bisphenol F (BPF), bisphenol S (BPS), and benzo[a]pyrene (BaP) stocks were provided by the Sherr laboratory and dissolved in DMSO. Stock concentrations were 4 × 10⁻³ M (BPA), 2 × 10⁻² M (BPF), 2 × 10⁻² M (BPS), and 10 mM (BaP). For treatments, bisphenols were added to culture media at final concentrations of 5 nM or 15 nM (1 µL per mL media), with DMSO serving as vehicle control. BaP was added at a final concentration of 1 µM (1 µL per 10 mL media) during specified exposure windows.

### Morphological analysis and developmental disruption index (DDI)

On day 14 post-plating, full-gel images were acquired using a Nikon confocal microscope. Image filenames were blinded prior to manual quantification. Four hydrogels per patient per condition were analyzed for: (i) total number of structures per gel, (ii) number of structures containing lobules, and (iii) presence of abnormal morphologies. Abnormal structures were defined as organoids exhibiting either (a) clustered lobules (≥3 lobules in a localized region rather than evenly distributed) or (b) lobules emerging directly from the primary duct rather than secondary or tertiary branches (“Brussels sprout” phenotype). For each patient, measurements from four gels were averaged and normalized to match DMSO controls. A composite developmental disruption index (DDI) was calculated using three criteria: Structure number: 0 = no change; 1 = 1.5–1.9-fold change; 2 = ≥2-fold change vs. DMSO; Percentage of structures with lobules: 0 = <2-fold; 1 = 2–4.9-fold; 2 = ≥5-fold vs. DMSO; Abnormal morphology: binary (0 = absent; 1 = present).

The DDI score for each condition and patient was calculated as the mean of these three metrics. For visualization, each patient was assigned a unique symbol. Normality was assessed using the Kolmogorov–Smirnov test. As data were not normally distributed, statistical comparisons were performed using the Wilcoxon signed-rank test (*p* ≤ 0.05).

### Live-cell tracking

On day 13, hydrogels were stained with Cytopainter Cell Tracking Dye (Abcam, ab138891) diluted 1:500 in complete MEGM media and incubated for 30 min at 37 °C with 5% CO₂. Gels were washed twice with fresh media prior to imaging.

### Imaging and AI-assisted segmentation

Brightfield and GFP images were acquired on day 13 using a Nikon confocal microscope. Five z-stacks per gel were collected and combined into maximum intensity projections. For segmentation model development, >10 full-gel images underwent manual annotation to generate binary masks identifying individual organoids. Annotated images were used to train a segmentation model using the Segment.ai function in Nikon NIS-Elements software. Following training, the model was applied to new images to extract organoid-level metrics including area, count, and circularity. Circularity was calculated as: 4π × (Area / Perimeter²)

### Bulk RNA sequencing

#### Sample preparation and sequencing

Six patient-derived organoid cultures were established as described above. For each patient, cells were seeded into four hydrogels per condition (4,000 cells per gel) and cultured for 21 days. Hydrogels were flash frozen and stored at −80 °C. RNA was extracted using TRIzol (Thermo Fisher Scientific, 15596018) followed by purification with the RNeasy Mini Kit (Qiagen, 74106). RNA quality and concentration were assessed using a NanoDrop 2000. Library preparation and paired-end sequencing were performed by Novogene using poly(A) enrichment and strand-specific library construction. Sequencing was conducted on an Illumina NovaSeq X-Plus platform.

#### Quality control and analysis

Reads were processed using fastp to remove adapters and low-quality reads. Alignment was performed with HISAT2 (v2.2.1), transcript assembly with StringTie (v2.2.3), and read counting with featureCounts (v2.0.6). Differential expression analysis was conducted using DESeq2 (v1.42.0), with adjusted *p*-values calculated via the Benjamini–Hochberg method. Genes with adjusted *p* ≤ 0.05 and |log₂ fold change| > 1 were retained for downstream analyses. GO and GSEA analyses were conducted using clusterProfiler (v4.8.1). Protein–protein interaction networks were generated using Cytoscape (v3.10.4) with STRING-derived interaction scores. Cell population deconvolution was performed using CIBERSORTx with a human breast scRNA-seq reference (Gray *et al.*, 2022). EMT-specific GSEA analyses were performed using cluster gene sets from ^28^.

#### Differential expression analysis and derivation of the 3BPX exposure signature

Raw gene-level counts from organoids exposed to 3BPX or DMSO were analyzed using Ensembl gene identifiers. Donor-matched differential expression analysis was performed with DESeq2^47^ using a design that accounted for donor and treatment condition, comparing 3BPX-treated and DMSO control samples within donor pairs. Genes with fewer than 10 total counts across all samples were excluded. Wald statistics were used for inference, and p values were adjusted for multiple testing using the Benjamini–Hochberg method.

Ensembl gene identifiers were subsequently mapped to gene symbols. Where multiple Ensembl features mapped to the same gene symbol, the feature with the smallest adjusted p value was retained. The 3BPX exposure signature was then defined as two non-overlapping gene sets: an upregulated set consisting of genes with adjusted P < 0.05 and log₂ fold change > 0, and a downregulated set consisting of genes with adjusted P < 0.05 and log₂ fold change < 0.

#### Projection of the 3BPX transcriptional signature into bulk breast cancer cohorts

To assess whether the 3BPX transcriptional program was represented in human breast tumors, the organoid-derived signature was projected into bulk expression datasets from TCGA-BRCA, METABRIC, and SCAN-B. In each cohort, gene symbols were standardized to uppercase and intersected with the 3BPX signature gene sets before scoring. Signature enrichment was quantified using GSVA^48^, and a composite 3BPX score was defined for each sample as the enrichment score of the upregulated signature minus that of the downregulated signature.

For TCGA-BRCA **<u>[The Cancer Genome Atlas Network. Comprehensive molecular portraits</u> <u>of human breast tumours. Nature 490, 61–70 (2012).]</u>**, RNA-seq data from 1,095 primary breast tumors were analyzed using an internally curated expression object. Expression values were converted from Ensembl identifiers to gene symbols, and duplicate mappings were resolved by retaining the feature with the highest mean expression. PAM50 subtype information, histologic classification, and ER status were integrated for downstream analyses; ER status was obtained from the UCSC Xena **<u>[Goldman MJ et al. Visualizing and interpreting cancer</u> <u>genomics data via the Xena platform. Nat Biotechnol 38, 675–678 (2020).]</u>** TCGA BRCA clinical resource and merged at the patient level.

For METABRIC **[Curtis C et al. The genomic and transcriptomic architecture of 2,000 breast tumours reveals novel subgroups. Nature 486, 346–352 (2012).]**, log-transformed expression and clinical annotation data were analyzed from a curated breast cancer expression cohort. Expression values were already provided at the gene-symbol level. ER status and intrinsic subtype information were harmonized for downstream analyses. Histologic subtype information was not available in the curated object used here, and analyses in this cohort were therefore restricted to ER- and subtype-based comparisons.

For SCAN-B **[Saal LH et al. The Sweden Cancerome Analysis Network–Breast (SCAN-B) Initiative: a large-scale multicenter infrastructure towards implementation of breast cancer genomic analyses in the clinical routine. Genome Med 7, 20 (2015).]**, processed RNA-seq and clinical metadata were analyzed from a curated breast cancer expression cohort. When multiple expression profiles were available for the same case, values were averaged to derive one case-level profile before scoring. Histologic subtype, ER status, and PAM50 subtype were harmonized from available clinical annotations. Cases showing discordant PAM50 assignments across multiple profiles were flagged and excluded from subtype-based analyses.

Statistical comparisons of 3BPX scores between binary groups were performed using two-sided Wilcoxon rank-sum tests. Histology-specific differences in TCGA and SCAN-B were examined both overall and within ER-positive tumors. Associations with PAM50 subtype were tested using Kruskal–Wallis tests followed by pairwise Wilcoxon tests with Benjamini–Hochberg correction. In TCGA and SCAN-B, histology associations were further evaluated using linear models with 3BPX score as the outcome and histology plus PAM50 subtype as predictors, with ER status added in sensitivity analyses. Within-subtype comparisons of IDC and ILC were restricted to strata containing at least 20 samples of each histologic class.

### Single-cell atlas projection

Differential expression signatures were projected onto the Human Breast Cancer Single Cell Atlas^33^ using the cellxgene platform. Enrichment scores were computed separately for epithelial and stromal compartments. For the epithelial subpopulation showing strongest overlap with the 3BPX signature (Luminal B), defining gene sets were extracted and subjected to Hallmark pathway analysis using clusterProfiler (*p* < 0.05).

### Reduced representation bisulfite sequencing (RRBS)

RRBS was performed on pooled DNA from 6 (3BPX) or 10 (BaP ± bisphenol) patient-derived organoid cultures harvested at day 21. DNA extraction was performed using the QIAamp DNA Mini Kit (Qiagen, 51304). Libraries were prepared by CD Genomics using MspI digestion, adapter ligation, size selection (∼150 bp), bisulfite conversion (>99%), and PCR amplification. Sequencing was conducted on an Illumina HiSeq X Ten. Reads were processed with FastQC and aligned using Bismark and HISAT2. Differentially methylated regions (DMRs) were identified using Metilene with Mann–Whitney U and 2D Kolmogorov–Smirnov tests. A composite rank score (70% statistical significance, 30% effect size) was calculated, and the top 10% of ranked DMRs were retained for downstream analyses. Circos plots were generated using the circlize R package (v0.4.11).

#### Methylation panel construction

For downstream methylation projection analyses, the retained top-ranked DMRs from the organoid RRBS comparisons DMSO vs BaP, DMSO vs 3BPX, DMSO+BaP vs BPA+BaP, DMSO+BaP vs BPF+BaP, and DMSO+BaP vs BPS+BaP were carried forward. DMRs were classified using the CD Genomics-provided genomic annotation columns: regions annotated to promoter, exon, or gene body were considered gene-associated, whereas regions lacking all three annotations were considered intergenic. Within each comparison, direction, and annotation class, DMRs with at least 5 CpGs were ranked by q-value, with ties resolved by absolute methylation difference and CpG count, and the top 10% were retained for downstream scoring.

#### Projection to TCGA-BRCA methylation data and statistical analysis

Public Illumina HumanMethylation450 methylation data from TCGA-BRCA^49^ were analyzed using an internally curated hg38-aligned TCGA-BRCA methylation resource containing primary tumors and adjacent normal breast samples. To avoid duplicate representation of tumors, one primary tumor sample per patient was retained, prioritizing aliquot 01A over 01B when both were available, while all adjacent normal samples were kept. Clinical metadata were assembled from the curated object, including PAM50 subtype and histology. ER status was obtained from the UCSC Xena^50^ TCGA BRCA clinical matrix, harmonized to ER+ or ER−, and merged by 12-character TCGA patient identifier.

Projection was performed at the DMR level. RRBS DMR intervals were expanded by ±500 bp and intersected with TCGA 450K probes in matched hg38 coordinates. For each DMR with at least one overlapping probe, probe-level beta values were averaged within sample to obtain a DMR-level methylation value. These DMR-level values were then z-scored across the TCGA analysis cohort. To preserve the expected organoid directionality, scores from hypomethylated RRBS DMRs were multiplied by −1, whereas hypermethylated DMRs retained a positive sign. For each sample, the final panel score was calculated as the mean of these direction-aligned DMR-level z-scores across all usable DMRs in the panel. A panel was considered measurable if it contained at least 5 measurable hypermethylated DMRs or at least 5 measurable hypomethylated DMRs.

Tumor-versus-normal comparisons of panel scores were performed using two-sided Wilcoxon rank-sum tests, and effect sizes were summarized using Cliff’s delta^51^. PAM50 subtype associations were evaluated using Kruskal–Wallis tests followed by pairwise Wilcoxon tests with Benjamini–Hochberg correction; ε² was used as a non-parametric effect size for subtype-level comparisons. Histology-specific differences were first examined in tumor samples using two-sided Wilcoxon rank-sum tests comparing IDC and ILC overall, and separately within ER-positive tumors; effect sizes were summarized using Cliff’s delta, and p-values were adjusted using the Benjamini–Hochberg method across panels. Histology-specific associations were then further evaluated in tumor samples using linear models with panel score as the outcome and histology plus PAM50 subtype as predictors, with ER status added in sensitivity analyses.

## Data availability

Primary RNAseq and RRBS data (fastq files) and raw-methylation calls are deposited to the Gene Expression Omnibus (GEO) repository under study accession numbers:

## Acknowledgements

We thank the members of the FTC Consortium for helpful discussions and David Sherr for critical evaluation of the manuscript. We gratefully acknowledge Karla Murga, Daniela Requena, and Megan Maloney at Tufts Biomedical Repository for tissue support. This research was supported by Find The Cause (FTC) Breast Cancer Foundation (to C.K. and S.M.).

## Author contributions

M.P., M.S, and C.K. conceived the project and designed experiments. M.P., M.S. performed experiments. M.P., H.N. and A.C performed sequencing analysis RNA-seq and RRBs data analysis and interpretation S.M. M.P. and C.K. wrote the manuscript.

## Supplemental Figure Legends

**Supplemental Figure 1. Single-cell–derived human breast organoid development and AI-assisted morphometric analysis. (A)** Schematic illustrating the single-cell–derived breast organoid model, showing progressive stages of organoid morphogenesis over time from an individual patient sample. Representative bright-field images depict structural maturation from early ductal formation through lobule-like architectures across developmental time points. **(B)** Workflow for development of the AI-assisted organoid segmentation pipeline. Organoids embedded within hydrogels were manually annotated across >10 training images, used to train a segmentation model, and subsequently applied to new images. Representative examples of manual annotation and AI-based segmentation are shown.

**(C)** Representative bright-field images illustrating organoid morphology following exposure to individual endocrine-disrupting chemicals (EDCs), with four representative organoids shown per treatment condition (n = 4). **(D)** Representative images demonstrating organoid morphological complexity stratified by circularity measurements, illustrating classification into high, moderate, low, or absent complexity states.

**Supplemental Figure 2. Dose-dependent morphological disruption and pathway enrichment following bisphenol exposure in human breast organoids. (A)** Representative bright-field microscopy images of organoids exposed to individual bisphenols (5 nM) at day 14 (n = 3 patient donors) (left), with corresponding quantitative assessments of lobule number, total number of structures, and abnormal structure ratio (right)**. (B)** Representative bright-field microscopy images of organoids exposed to individual bisphenols (15 nM) at day 14 (n = 3 patient donors), with corresponding quantitative morphometric analyses (right). **(C)** Developmental disruption index (DDI) scores for organoids exposed to bisphenols at 15 nM (left) and 5 nM (right) concentrations at day 14. **(A–C)** Each symbol represents an individual patient-derived organoid; symbol shapes correspond to patient identity. Statistical significance was assessed using the Wilcoxon signed-rank test. **(D)** Gene sets corresponding to four clusters identified by Cytoscape network analysis were subjected to over-representation analysis (ORA) using Hallmark and Gene Ontology (GO) pathway libraries. Shown are the raw Hallmark ORA outputs for each cluster; significantly enriched pathways are outlined in red (adjusted *p* < 0.05).

**Supplemental Figure 3. BaP exposure induces dose-dependent cell death and DNA damage in developing human breast organoids. (A)** Representative fluorescence images of organoids exposed to benzo[a]pyrene (BaP) for 48 hours at either 1 µM or 10 µM during developmental days 5–7, followed by continued culture to day 14. Organoids were stained with Zombie Green (1:1000) to assess cell death and immunostained for γ-H2AX (1:1000; red) to detect DNA damage. **(B)** Gene Ontology (GO) biological process enrichment analysis of shared differentially expressed genes in response to BaP or 3BPX exposure.

**Supplemental Figure 4. Quantitative assessment of morphological disruption following combined bisphenol and BaP exposure. (A)** Quantitative morphometric analyses of organoids (n = 10 patient donors) exposed to individual bisphenols at 5 nM or 15 nM in combination with benzo[a]pyrene (BaP; 1 µM). Bisphenol exposure was continuous throughout organoid development at the indicated concentrations, while BaP was administered for 48 hours during developmental days 5–7. Morphological quantifications were performed on day 14. Statistical significance was assessed using the Wilcoxon signed-rank test. **(B)** Developmental disruption index (DDI) scores for organoids exposed to bisphenols in the presence or absence of BaP at day 14. **(A,B)** Each data point represents an individual patient-derived organoid; symbol shapes denote patient identity.

**Supplemental Figure 5. Additional analyses supporting subtype and histology associations of exposure-derived DNA methylation signatures in TCGA-BRCA. (A)** Distribution of RRBS-derived methylation panel scores across PAM50 breast cancer subtypes in TCGA-BRCA. The BaP+BPS-associated signature (B_vs_E) exhibits the strongest subtype stratification, with higher scores in Basal tumors relative to Luminal and Normal-like subtypes. The 3BPX-derived signature (A_vs_F) also shows significant subtype structure, with enrichment in Normal-like and Basal tumors compared to Luminal subtypes. Other panels (BaP+BPA (B_vs_C), BaP+BPF (B_vs_D) display weaker but statistically significant subtype associations, while A_vs_B shows no meaningful subtype structure. **(B) C**omparison of exposure-derived methylation panel scores between invasive lobular carcinoma (ILC) and invasive ductal carcinoma (IDC) across all tumors (IDC = 328, ILC = 116). The 3BPX-derived methylation signature (A_vs_F) is significantly elevated in ILC relative to IDC, while BaP-associated co-exposure signatures (BaP+BPS (B_vs_E), BaP+BPA (B_vs_C) are reduced in ILC. No significant differences are observed for BaP (A_vs_B) or BaP+BPF (B_vs_D). **(C)** Histology comparison restricted to ER⁺ tumors (ER⁺ IDC = 213, ER⁺ ILC = 109). The enrichment of the 3BPX-derived methylation signature (A_vs_F) in ILC persists within ER⁺ disease, indicating that this association is independent of ER status. A weaker inverse association is observed for BaP+BPA (B_vs_C), while other panels show no significant or only borderline differences.

## Supplemental Table Legends

**Supplemental Table 1: 3D-HCS summary of organoid phenotypes following exposure to individual chemicals**. Reported outcomes include changes in total organoid number, shifts in organoid size distribution (extra-small, small, medium, large, extra-large), and alterations in structural complexity categories (no, low, moderate, high complexity).

**Supplemental Table 3: Patient and clinical characteristics of breast tissue specimens used in this study**. Dashes indicate information that was unavailable or not reported. Pregnancy history is shown as the number of prior pregnancies when available. All tissue samples wer**e** collected under the Institutional Review Board–approved protocols with informed consent.

**Supplemental Table 4:Genes included in protein–protein interaction (PPI) network analysis of differentially expressed genes.** Differentially expressed genes identified from 3BPX RNA-seq analysis used to construct protein–protein interaction (PPI) networks in Cytoscape using STRING-derived interaction data (confidence score 400–700). Genes are listed according to their inclusion in upregulated and downregulated networks corresponding to Figure 2E & 2H. These gene sets were used to identify highly connected hub genes and to define major functional modules within each network.

**Supplemental Table 5. Gene lists corresponding to stromal clusters identified by scRNA-seq across human breast tumors.** Stromal populations were resolved into 14 distinct clusters, six of which corresponding to previously described cancer-associated fibroblast (CAF) states and myofibroblast populations. These gene signatures were used to annotate stromal cell identities and to support downstream analyses of fibroblast heterogeneity.

## References

1 Ye, X. et al. Urinary Concentrations of Bisphenol A and Three Other Bisphenols in Convenience Samples of U.S. Adults during 2000-2014. Environ Sci Technol 49, 11834–11839 (2015). 10.1021/acs.est.5b02135

2 Vandenberg, L. N. et al. Urinary, circulating, and tissue biomonitoring studies indicate widespread exposure to bisphenol A. Environ Health Perspect 118, 1055–1070 (2010). 10.1289/ehp.0901716

3 Stanfield, Z. et al. Characterizing Chemical Exposure Trends from NHANES Urinary Biomonitoring Data. Environ Health Perspect 132, 17009 (2024). 10.1289/ehp12188

4 Clayton, E. M., Todd, M., Dowd, J. B. & Aiello, A. E. The impact of bisphenol A and triclosan on immune parameters in the U.S. population, NHANES 2003-2006. Environ Health Perspect 119, 390–396 (2011). 10.1289/ehp.1002883

5 Young, A. S. et al. Hormone receptor activities of complex mixtures of known and suspect chemicals in personal silicone wristband samplers worn in office buildings. Chemosphere 315, 137705 (2023). 10.1016/j.chemosphere.2022.137705

6 Samon, S. et al. Measuring semi-volatile organic compound exposures during pregnancy using silicone wristbands. Chemosphere 339, 139778 (2023). 10.1016/j.chemosphere.2023.139778

7 Macon, M. B. & Fenton, S. E. Endocrine disruptors and the breast: early life effects and later life disease. J Mammary Gland Biol Neoplasia 18, 43–61 (2013). 10.1007/s10911-013-9275-7

8 Yang, Y. et al. Per- and Polyfluoroalkyl Substances (PFAS) in Consumer Products: An Overview of the Occurrence, Migration, and Exposure Assessment. Molecules 30 (2025). 10.3390/molecules30050994

9 Parrish, M., Traugh, N., Seraj, M. & Kuperwasser, C. Field cancerization, accelerated aging, and immunosuppression: the rapid rise of hormone-sensitive and early-onset breast cancer. NPJ Breast Cancer 11, 128 (2025). 10.1038/s41523-025-00840-w

10. Institute of Medicine Committee on the Implications of Dioxin in the Food, S. in Dioxins and Dioxin-like Compounds in the Food Supply: Strategies to Decrease Exposure (National Academies Press (US) Copyright 2003 by the National Academy of Sciences. All rights reserved., 2003).

11. National Research Council Committee on Hormonally Active Agents in the, E. in Hormonally Active Agents in the Environment (National Academies Press (US) Copyright 1999 by the National Academy of Sciences. All rights reserved., 1999).

12 Soto, A. M. & Sonnenschein, C. Environmental causes of cancer: endocrine disruptors as carcinogens. Nat Rev Endocrinol 6, 363–370 (2010). 10.1038/nrendo.2010.87

13 Habert, R. et al. Concerns about the widespread use of rodent models for human risk assessments of endocrine disruptors. Reproduction 147, R119–129 (2014). 10.1530/rep-13-0497

14 Lacouture, A. et al. Estrogens and endocrine-disrupting chemicals differentially impact the bioenergetic fluxes of mammary epithelial cells in two- and three-dimensional models. Environ Int 179, 108132 (2023). 10.1016/j.envint.2023.108132

15 Alonso-Magdalena, P. et al. Bisphenol-A acts as a potent estrogen via non-classical estrogen triggered pathways. Mol Cell Endocrinol 355, 201–207 (2012). 10.1016/j.mce.2011.12.012

16 Golden, R., Gandy, J. & Vollmer, G. A review of the endocrine activity of parabens and implications for potential risks to human health. Crit Rev Toxicol 35, 435–458 (2005). 10.1080/10408440490920104

17 Morito, K. et al. Interaction of phytoestrogens with estrogen receptors alpha and beta. Biol Pharm Bull 24, 351–356 (2001). 10.1248/bpb.24.351

18 Gaillard, L., Barouki, R., Blanc, E., Coumoul, X. & Andréau, K. Per- and polyfluoroalkyl substances as persistent pollutants with metabolic and endocrine-disrupting impacts. Trends Endocrinol Metab 36, 249–261 (2025). 10.1016/j.tem.2024.07.021

19 Mándi, Y. & Vécsei, L. The kynurenine system and immunoregulation. J Neural Transm (Vienna) 119, 197–209 (2012). 10.1007/s00702-011-0681-y

20 Gelboin, H. V. Benzo[alpha]pyrene metabolism, activation and carcinogenesis: role and regulation of mixed-function oxidases and related enzymes. Physiol Rev 60, 1107–1166 (1980). 10.1152/physrev.1980.60.4.1107

21 Kortenkamp, A. Ten years of mixing cocktails: a review of combination effects of endocrine-disrupting chemicals. Environ Health Perspect 115 Suppl 1, 98–105 (2007). 10.1289/ehp.9357

22 Rauner, G., Gupta, P. B. & Kuperwasser, C. From 2D to 3D and beyond: the evolution and impact of in vitro tumor models in cancer research. Nat Methods 22, 1776–1787 (2025). 10.1038/s41592-025-02769-1

23 Rauner, G. et al. Breast tissue regeneration is driven by cell-matrix interactions coordinating multi-lineage stem cell differentiation through DDR1. Nat Commun 12, 7116 (2021). 10.1038/s41467-021-27401-6

24 Rauner, G. et al. Single-cell organogenesis captures complex breast tissue formation in three dimensions. Development 152 (2025). 10.1242/dev.204813

25 Trepicchio, C. et al. DDR1 regulates RUNX1-CBFβ to control breast stem cell differentiation. Stem Cell Reports 20, 102576 (2025). 10.1016/j.stemcr.2025.102576

26 Vandenberg, L. N. et al. Hormones and endocrine-disrupting chemicals: low-dose effects and nonmonotonic dose responses. Endocr Rev 33, 378–455 (2012). 10.1210/er.2011-1050

27 Steen, C. B., Liu, C. L., Alizadeh, A. A. & Newman, A. M. Profiling Cell Type Abundance and Expression in Bulk Tissues with CIBERSORTx. Methods Mol Biol 2117, 135–157 (2020). 10.1007/978-1-0716-0301-7_7

28 Liang, L. et al. Meta-Analysis of EMT Datasets Reveals Different Types of EMT. PLoS One 11, e0156839 (2016). 10.1371/journal.pone.0156839

29 Chang, Y. et al. Comparative mechanisms of PAH toxicity by benzo[a]pyrene and dibenzo[def,p]chrysene in primary human bronchial epithelial cells cultured at air-liquid interface. Toxicol Appl Pharmacol 379, 114644 (2019). 10.1016/j.taap.2019.114644

30 Caporale, N. et al. From cohorts to molecules: Adverse impacts of endocrine disrupting mixtures. Science 375, eabe8244 (2022). 10.1126/science.abe8244

31 Barroso-Sousa, R. & Metzger-Filho, O. Differences between invasive lobular and invasive ductal carcinoma of the breast: results and therapeutic implications. Ther Adv Med Oncol 8, 261–266 (2016). 10.1177/1758834016644156

32 Lee, A. H., Happerfield, L. C., Millis, R. R. & Bobrow, L. G. Inflammatory infiltrate in invasive lobular and ductal carcinoma of the breast. Br J Cancer 74, 796–801 (1996). 10.1038/bjc.1996.438

33 Chen, A., Kroehling, L., Ennis, C. S., Denis, G. V. & Monti, S. A highly resolved integrated transcriptomic atlas of human breast cancers. NARGAB (2026). 10.1093/nargab/lqaf217

34 Gao, Y. et al. Cross-tissue human fibroblast atlas reveals myofibroblast subtypes with distinct roles in immune modulation. Cancer Cell 42, 1764–1783.e1710 (2024). 10.1016/j.ccell.2024.08.020

35 Du, Y. et al. Integration of Pan-Cancer Single-Cell and Spatial Transcriptomics Reveals Stromal Cell Features and Therapeutic Targets in Tumor Microenvironment. Cancer Res 84, 192–210 (2024). 10.1158/0008-5472.Can-23-1418

36 Croizer, H. et al. Deciphering the spatial landscape and plasticity of immunosuppressive fibroblasts in breast cancer. Nat Commun 15, 2806 (2024). 10.1038/s41467-024-47068-z

37 Kieffer, Y. et al. Single-Cell Analysis Reveals Fibroblast Clusters Linked to Immunotherapy Resistance in Cancer. Cancer Discov 10, 1330–1351 (2020). 10.1158/2159-8290.Cd-19-1384

38 Gómez-Cuadrado, L. et al. Characterisation of the Stromal Microenvironment in Lobular Breast Cancer. Cancers (Basel) 14 (2022). 10.3390/cancers14040904

39 Ames, B. N., McCann, J. & Yamasaki, E. Methods for detecting carcinogens and mutagens with the Salmonella/mammalian-microsome mutagenicity test. Mutat Res 31, 347–364 (1975). 10.1016/0165-1161(75)90046-1

40 Hanahan, D. Hallmarks of Cancer: New Dimensions. Cancer Discov 12, 31–46 (2022). 10.1158/2159-8290.Cd-21-1059

41 Desgraupes, S., Etienne, L. & Arhel, N. J. RANBP2 evolution and human disease. FEBS Lett 597, 2519–2533 (2023). 10.1002/1873-3468.14749

42 Ellington, T. D. et al. Trends in Breast Cancer Incidence, by Race, Ethnicity, and Age Among Women Aged ≥20 Years - United States, 1999-2018. MMWR Morb Mortal Wkly Rep 71, 43–47 (2022). 10.15585/mmwr.mm7102a2

43 Giaquinto, A. N., Freedman, R. A., Newman, L. A., Jemal, A. & Siegel, R. L. Lobular breast cancer statistics, 2025. Cancer 131, e70061 (2025). 10.1002/cncr.70061

44 Heaphy CM, Griffith JK, Bisoffi M. Mammary field cancerization: molecular evidence and clinical importance. Breast Cancer Res Treat. 118, 229–39 (2009). doi: 10.1007/s10549-009-0504-0.

45. Youssef, K.K., Narwade, N., Arcas, A. et al. Two distinct epithelial-to-mesenchymal transition programs control invasion and inflammation in segregated tumor cell populations. Nat Cancer 5, 1660–1680 (2024). 10.1038/s43018-024-00839-5

46. Aibar, S., González-Blas, C., Moerman, T. et al. SCENIC: single-cell regulatory network inference and clustering. Nat Methods 14, 1083–1086 (2017). 10.1038/nmeth.4463

47. Love MI, Huber W, Anders S. Moderated estimation of fold change and dispersion for RNA-seq data with DESeq2. Genome Biol 15, 550 (2014).

48. Hänzelmann S, Castelo R, Guinney J. GSVA: gene set variation analysis for microarray and RNA-seq data. BMC Bioinformatics 14, 7 (2013).

49. The Cancer Genome Atlas Network. Comprehensive molecular portraits of human breast tumours. Nature 490, 61–70 (2012).

50. Goldman MJ et al. Visualizing and interpreting cancer genomics data via the Xena platform. Nat Biotechnol 38, 675–678 (2020).

51. Cliff N. Ordinal Methods for Behavioral Data Analysis. Routledge, 1996.

