## Supplementary material for "Environmental chemical mixtures reprogram mammary epithelial development to epigenetic states associated with breast cancer": Supplmental Figures

### Supplmental Figure 1

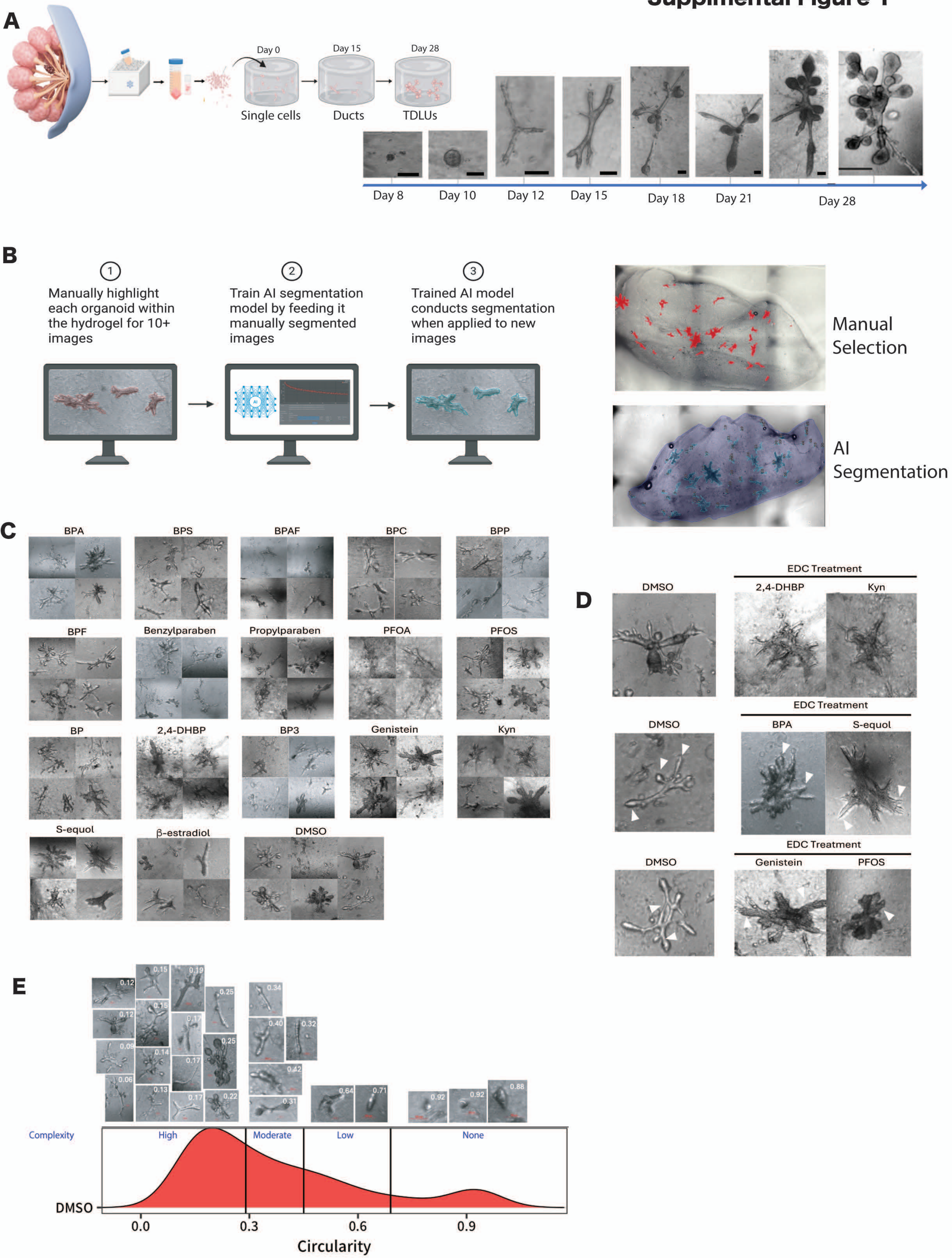

### Supplemental Figure 2

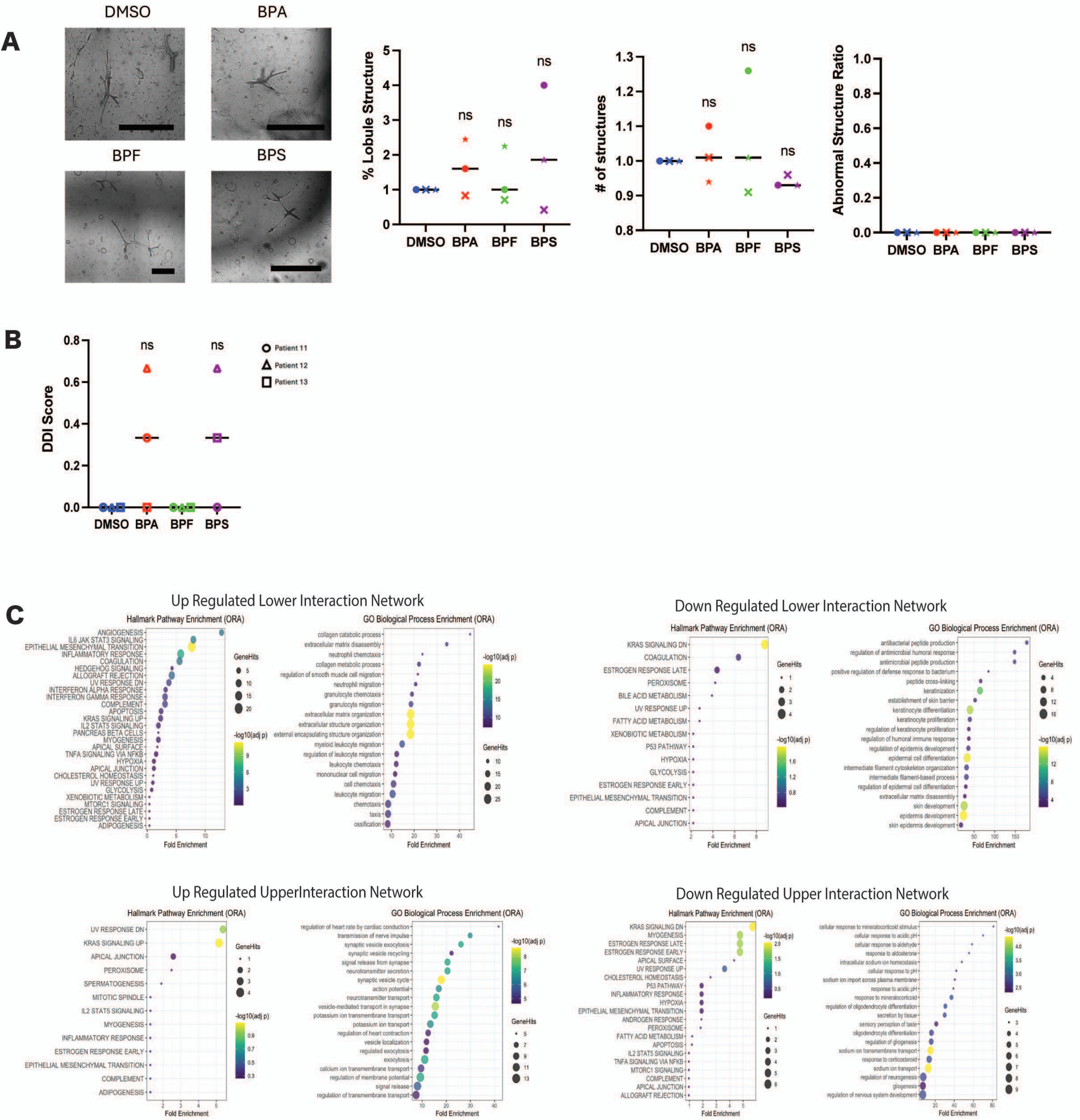

### Supplemental Figure 3

A

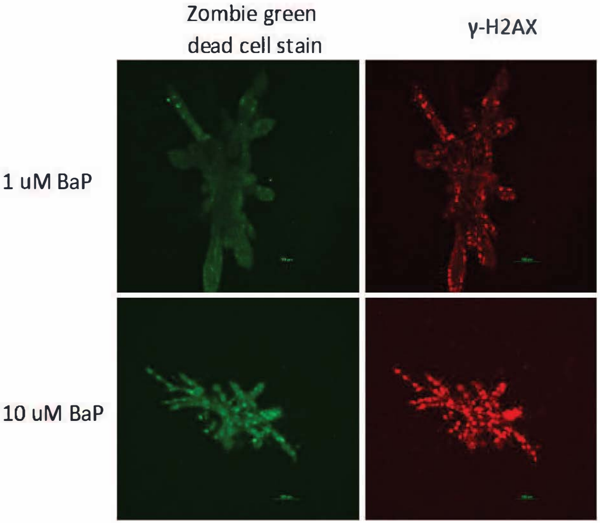


B

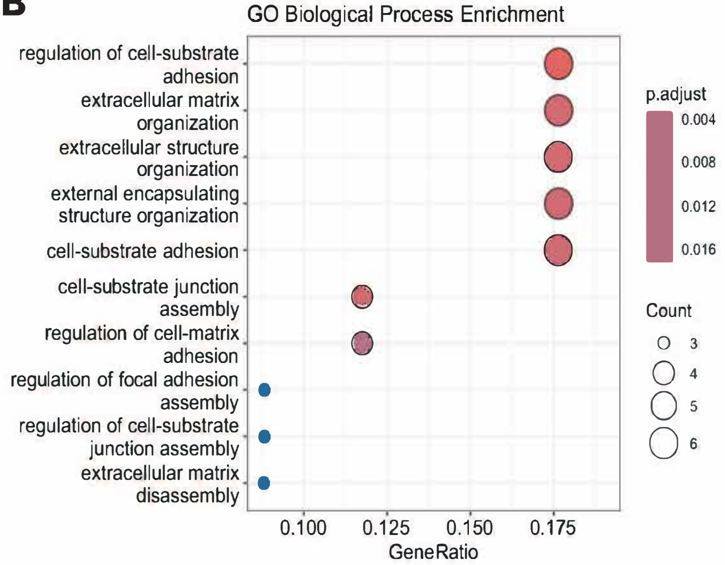

### Supplemental Figure 4

**A**

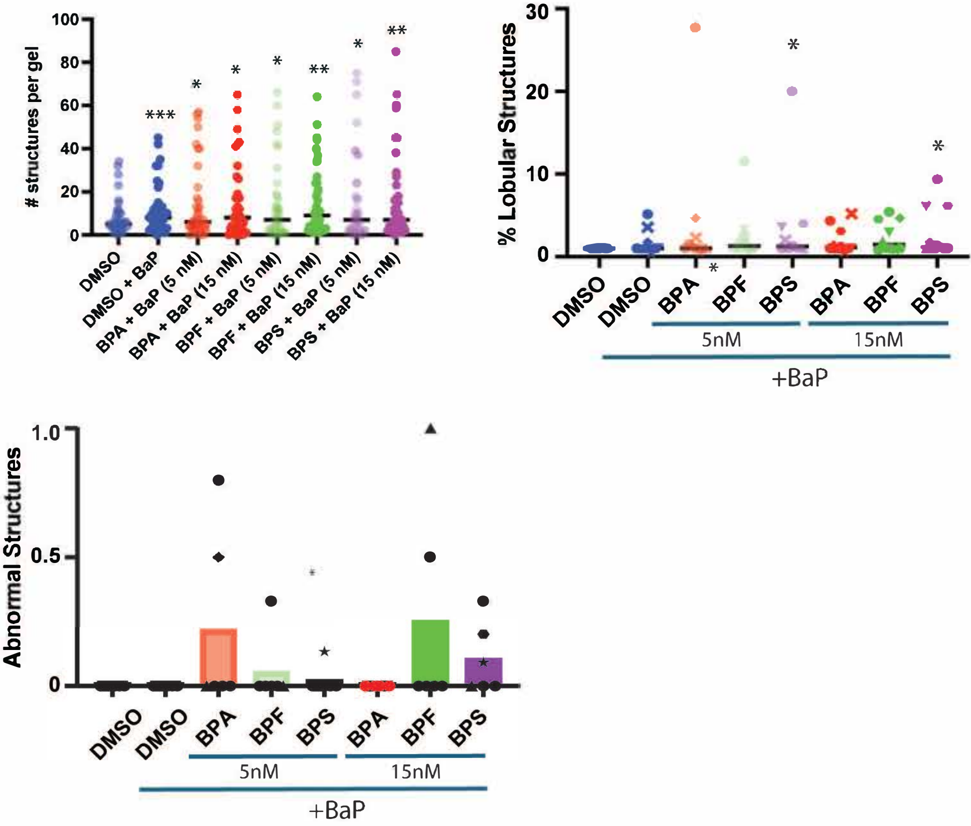


**B**

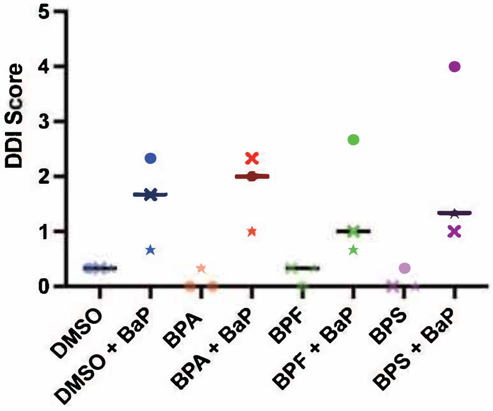

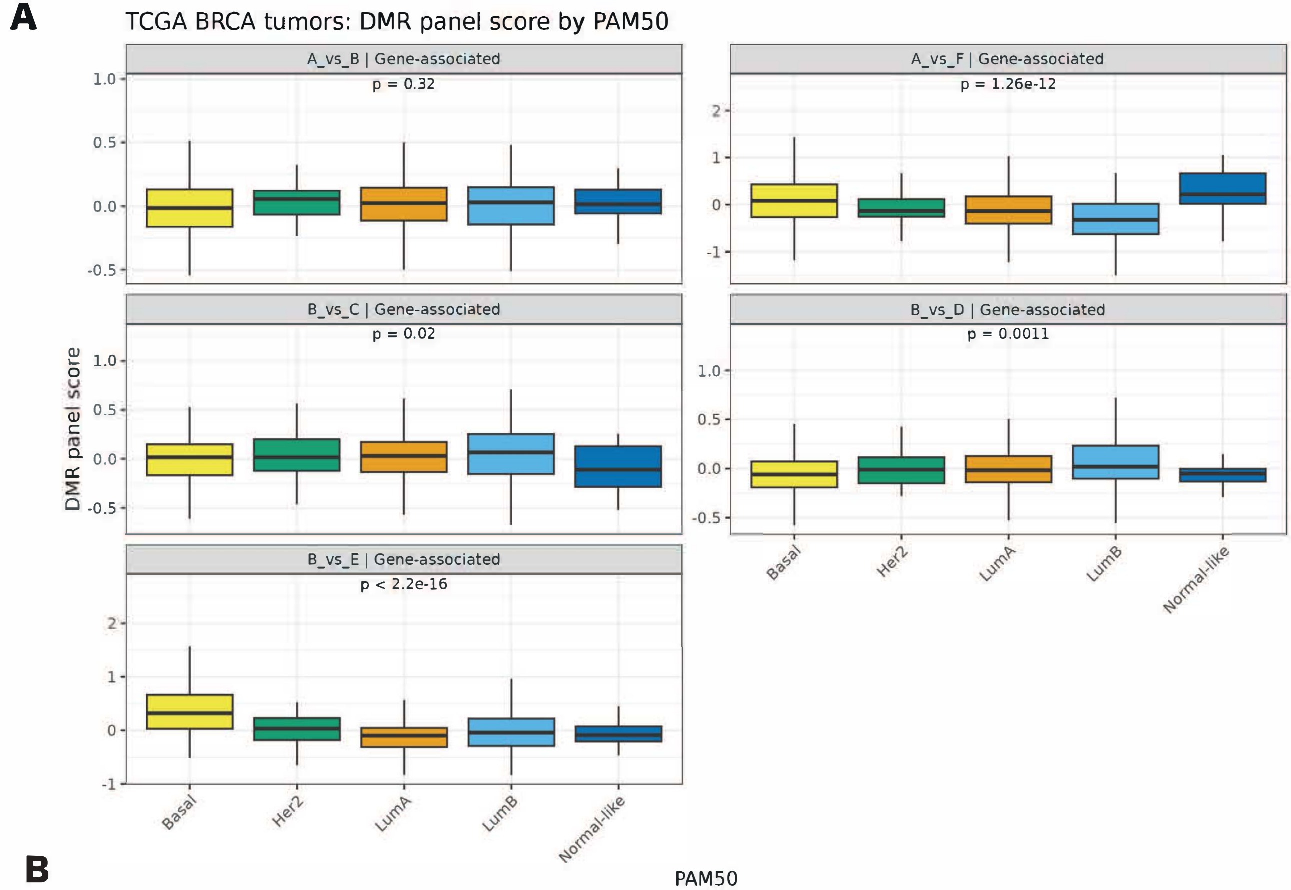


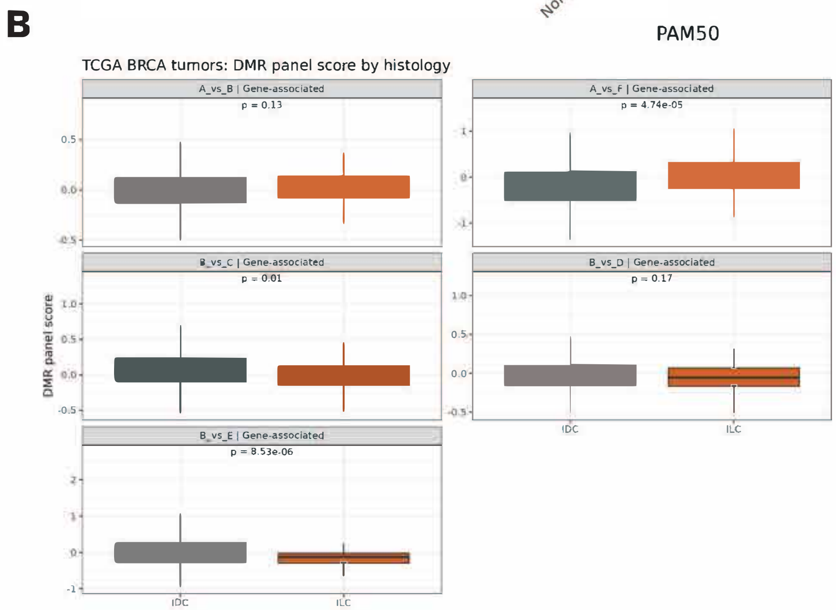


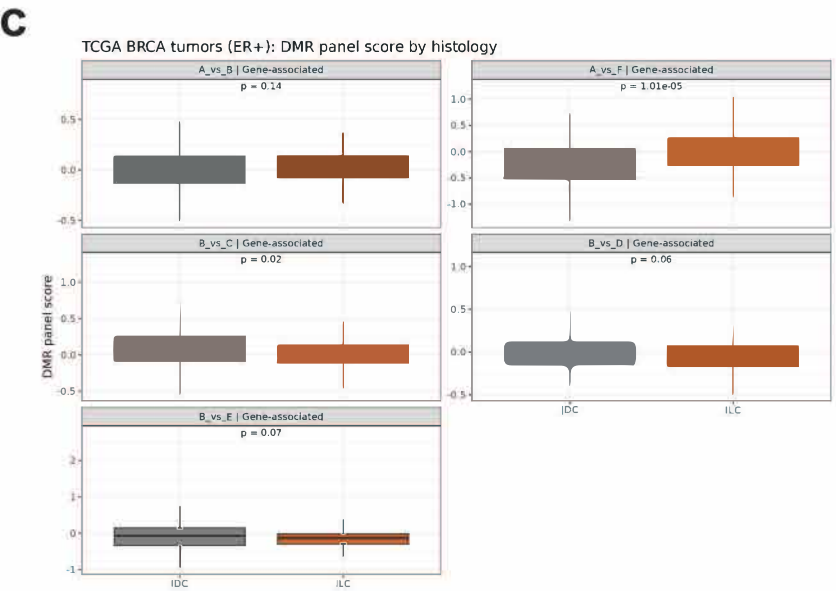


**Supplemental Figure 5**
